# High-throughput light-sheet 3D microscopy of whole rodent brain microstructure for integration with MRI

**DOI:** 10.64898/2026.10.12.758500

**Authors:** Jenna M. Hanmer, Francesco D’Antonio, Stephania Assimopoulos, Jose-Pedro Manzano-Patron, Sydney C. Wood, Mohamed Tachrount, Qijia Shen, Martin Craig, Cyril Lai, Graham J. Hickman, Edalat Radfar, Kim I. Chisholm, Tracy D. Farr, Alan Jasanoff, Anna Hoerder-Suabedissen, Zoltan Molnar, Clemence Ligneul, Cristiana Tisca, Saad Jbabdi, Amy F.D. Howard, Karla L. Miller, Elizabeth de Guzman, Jason P. Lerch, Stamatios N. Sotiropoulos

**Author notes:** These authors contributed equally (Francesco D’Antonio, Stephania Assimopoulos, Jose-Pedro Manzano-Patron, Elizabeth de Guzman, Jason P. Lerch, Stamatios N. Sotiropoulos).

## Abstract

Whole-brain cellular mapping is essential for understanding how differences in cellular architecture shape brain organisation and function. Magnetic Resonance Imaging (MRI) provides whole-brain coverage in humans and animal models but is only indirectly sensitive to cellular features. Integrating MRI with whole-brain microscopy is therefore critical for interpreting existing microstructural MRI measures and developing more specific ones. Here, we present a scalable cross-resolution framework integrating 3D light-sheet microscopy (LSM) of intact rodent brains with diffusion MRI (dMRI). By combining active tissue clearing and immunolabelling with accelerated LSM acquisitions, we achieve multiplexed whole-brain mapping of neuronal and glial populations on timescales comparable to conventional histology and 2D microscopy. This throughput enables LSM and MRI integration at scale, linking macroscopic MRI contrasts to spatially-matched microscopic cellular features within the same tissue. We apply this framework across eight genetically diverse mouse strains to develop predictive models of 3D microscopy-derived cellular features from dMRI contrasts. We demonstrate that data-driven combinations of state-of-the-art multi-shell dMRI measures predict neuronal and astrocytic features better than individual contrasts, across all animals. Our approach enables 3D microscopy to guide the development and interpretation of biologically-specific microstructural MRI measures, with the potential for translation to the living human brain.

## Introduction

Mapping cellular organisation across the whole brain is critical for understanding how neuronal and glial populations vary across individuals and how such differences shape brain architecture and function (Nowakowski et al., 2025). Beyond characterising normative variation, cellular brain maps are essential for uncovering disease mechanisms and the determinants of inter-individual variability in susceptibility, progression, severity, and recovery (Gabitto et al., 2024). However, achieving whole-brain cellular mapping remains a significant challenge because existing imaging approaches involve fundamental trade-offs between spatial coverage, resolution, biological specificity, and compatibility with *in vivo* applications.

Magnetic resonance imaging (MRI), and diffusion MRI (dMRI) in particular, provide the potential for 3D whole-brain measurements of tissue microstructure (Jelescu et al., 2022; Palombo et al., 2020; Zhang et al., 2012). MRI can be performed repeatedly, non-invasively and at high throughput, both *in vivo* and *ex vivo* (Miller et al., 2016; Tendler et al., 2022), making it uniquely suited to studying microstructural variation across the entire brain and in living humans. However, MRI operates at a relatively low spatial resolution and is only indirectly sensitive to cellular features, making the interpretation and biological specificity of MRI contrasts an open question. Optical microscopy (Antony et al., 2013; Balasubramanian et al., 2023) provides a complementary view, offering sub-micrometre resolution and high specificity when combined with molecular markers (Kronman et al., 2024; Lee et al., 2025; Wang et al., 2025) but approaches are inherently destructive and restricted to *ex vivo* tissue. Integrating MRI and whole-brain microscopy is therefore critical for interpreting existing microstructural MRI measures and developing new, more specific ones that are applicable *in vivo*.

Cross-scale, multimodal imaging offers a means of leveraging these complementary strengths by linking macroscopic MRI contrasts to microscopical cellular measurements. This integrative approach enables MRI methods to be calibrated, validated, and refined against high-resolution microscopy. Several studies have demonstrated the potential of this strategy by combining MRI with microscopy-derived atlases of the mouse brain (Han et al., 2026; Liang et al., 2022; Radhakrishnan et al., 2022), existing histological datasets (Wang et al., 2020), or conventional histology and 2D polarised light imaging in non-human primates (Calamante et al., 2012; Howard et al., 2023; Leergaard et al., 2010; Sepehrband et al., 2015). These studies provide important links between MRI measurements and underlying tissue architecture, however they rely on 2D microscopy and/or a limited number of thin tissue sections. This makes it difficult to capture the inherently 3D spatial heterogeneity of cellular organisation, to scale across whole brain volumes, and to determine whether relationships between MRI measures and cellular features generalise across brain regions, specimens and microstructural environments.

Three-dimensional fluorescence light-sheet microscopy (LSM) provides an opportunity to overcome these limitations. LSM enables volumetric imaging of samples rendered optically transparent via tissue clearing (Chung et al., 2013), providing cellular resolution across relatively large fields of view (Dodt et al., 2007; Huisken et al., 2004). When combined with immunolabelling for specific cellular markers (Yun et al., 2025), LSM can map cellular organisation and cytoarchitecture across both grey and white matter in 3D. Unlike 2D microscopy methods that require thin tissue sections, LSM can image intact specimens (Ueda et al., 2020). This preserves the 3D spatial organisation of the tissue, facilitates direct registration with inherently 3D MRI contrasts, and enables MRI–cellular relationships to be assessed continuously across anatomical structures and microstructural environments within the same brain.

Realising this potential at scale requires whole brain LSM workflows that provide uniform labelling, quantitative measurements, and sufficiently high throughput for integration with MRI, ideally across multiple specimens. However, tissue clearing, whole-brain immunolabeling, and LSM acquisition are inherently time-consuming and technically demanding (Weiss et al., 2021). Conventional clearing and immunolabeling approaches largely rely on *passive* diffusion of clearing agents and antibodies through tissue, which can be prohibitively slow, reducing throughput (Renier et al., 2016, 2014; Susaki et al., 2014; Tomer et al., 2014). Limited reagent penetration and probe depletion can also produce staining gradients, compromising the quantification of cellular features across the brain (Weiss et al., 2021). More recent *active* clearing and immunolabelling strategies use an external force, such as an electric field, to drive reagents and antibodies through tissue, accelerating processing and improving penetration uniformity (Cai et al., 2019; Kim et al., 2015; Lee et al., 2016). These approaches have begun to extend volumetric imaging from small tissue blocks (Stolp et al., 2018) to whole rodent brains (Johnson et al., 2023; Yau et al., 2023; Yun et al., 2025).

In this work, we present a scalable cross-resolution framework that integrates 3D LSM of intact rodent brains with dMRI to bridge cellular and macroscopic brain imaging. We further develop and optimise active electrotransport-based protocols to accelerate tissue clearing and immunolabelling, and achieve simultaneous whole-brain labelling of multiple cell types, including neurons and astrocytes, within two weeks. We pair these protocols with accelerated LSM acquisitions that enable whole-brain 3D cellular mapping at micrometre resolution in under five hours. Together, these advances achieve 3D cellular mapping on timescales comparable to conventional histology and 2D microscopy, providing the throughput required to integrate LSM with MRI at scale.

We apply this framework across eight genetically diverse mouse strains to investigate whether state-of-the-art multi-shell dMRI contrasts (Palombo et al., 2020) can predict underlying neuronal and astrocytic features across varied microstructural environments. This enables us to test not only the relationships between individual dMRI contrasts and microscopy-derived cellular features, but also whether data-driven combinations of complementary dMRI contrasts improve prediction of these cellular features. Furthering previous studies of MRI–3D microscopy integration (Gimenez et al., 2017; Goubran et al., 2019; Han et al., 2026; Johnson et al., 2023; Mansour et al., 2025; Stolp et al., 2018), our work brings together three key contributions: (i) accelerated processing and imaging of intact rodent brains, enabling whole-brain cellular mapping on timescales comparable to conventional histology; (ii) simultaneous multiplexed whole-brain labelling of multiple endogenous cellular targets using conventional antibodies that enables explorations of differential sensitivity of MRI markers to different cellular features; and (iii) a cross-scale analysis framework for directly relating 3D cellular maps with conventional (Jensen et al., 2005) and state-of-the-art microstructural dMRI measures (Palombo et al., 2020) acquired from the same specimens.

Taken together, these developments establish a scalable platform for using 3D microscopy to guide the development and interpretation of microstructural MRI measures. By directly linking macroscopic MRI contrasts to cell-type-specific measurements across the intact brain and across animals, we provide a principled approach for identifying combinations of MRI measurements that more specifically reflect underlying cellular architecture. Ultimately, this has the potential to improve the biological specificity of microstructural MRI measures that are translatable to the living human brain.

## Results

### A cross-scale workflow linking diffusion MRI to whole-brain 3D microscopy

We established a novel cross-scale imaging workflow that directly links multi-shell *ex vivo* diffusion MRI to multiplexed whole-brain 3D light-sheet microscopy in the same rodent brains (Fig 1A). This provides an integrated route from MRI-scale measurements of tissue microstructure to cellular-scale measurements of neuronal and glial organisation, enabling spatially matched comparisons across intact brains and avoiding the need of inferring MRI-cellular relationships from separate specimens or a limited number of tissue sections. To achieve this at scale, we developed and calibrated tissue processing and 3D microscopy protocols to preserve specimen integrity, achieve sufficient LSM throughput, and maintain compatibility with MRI.

**FIGURE 1.**
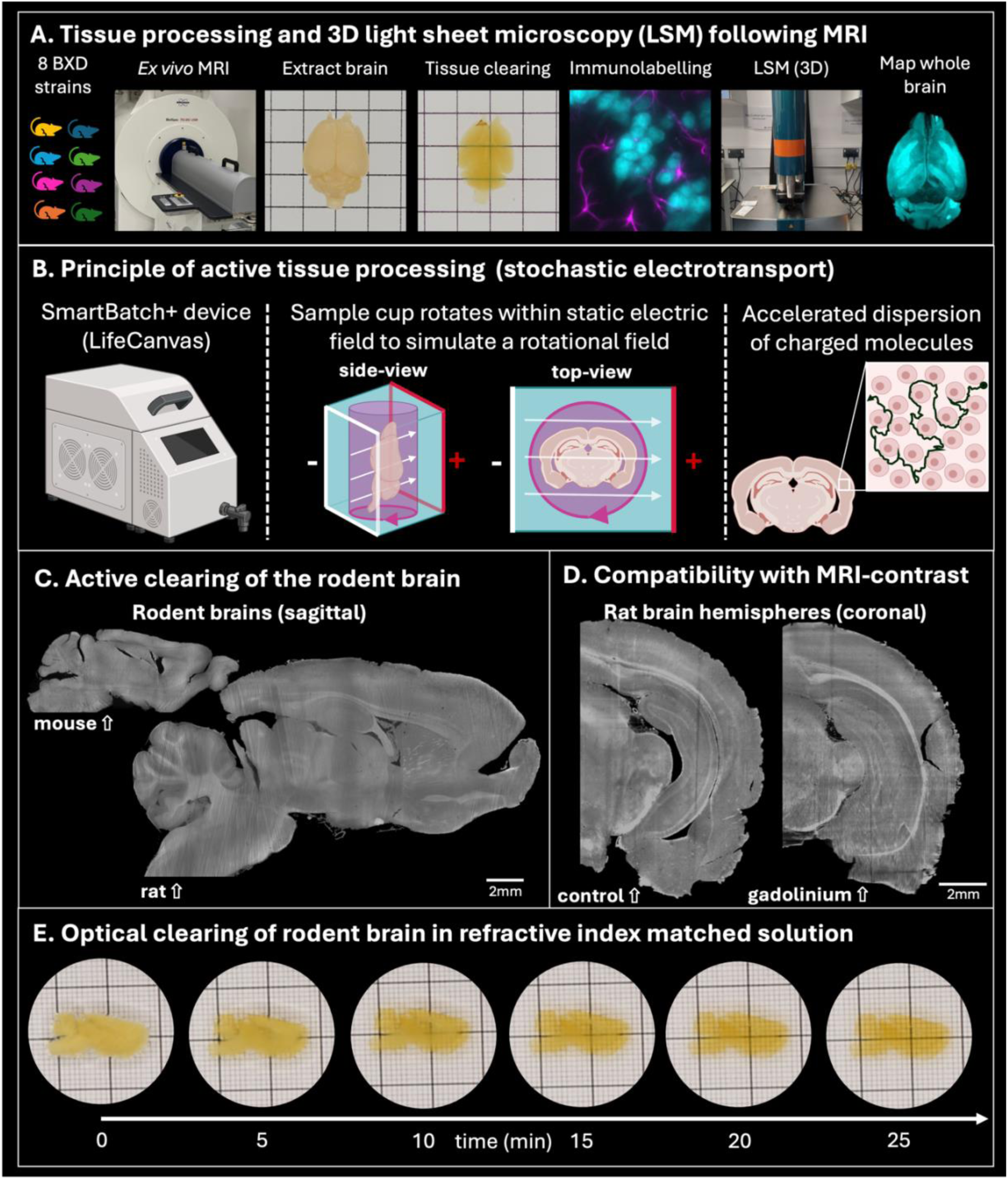
MRI, tissue clearing, immunolabelling, and light sheet microscopy on intact rodent brains. **(A)** Experimental workflow for high-resolution *ex vivo* MRI followed by tissue clearing, immunolabelling, refractive index matching and 3D light sheet microscopy of the same intact rodent brain (mouse). Grid scale 5mm. **(B)** Schematic of active tissue clearing and immunolabelling using the SmartBatch+ electrophoretic device. Left: The SmartBatch+ system, which accommodates up to 4 intact rat brains or 12 intact mouse brains. Middle: A rodent brain within the rotating sample holder (pink arrow) is exposed to a static electric field (white arrows) within the device’s inner chamber. Right: Rotation through the electric field promotes the uniform distribution of charged molecules, including antibodies, throughout dense tissue in a diffusion-like manner. **(C)** Background autofluorescence signal in the mouse (left) and rat (right) brains following active tissue clearing provides additional anatomical contrast. **(D)** Gadolinium-based MRI contrast agent can be included in sample preparation without compromising tissue clearing or subsequent light-sheet microscopy, as demonstrated in rat brain hemispheres. **(E)** Complete optical clearing, in index matched solution, of rat brain following MRI and tissue processing. Grid scale 1mm/1cm.

We applied this workflow to eight genetically distinct F1 hybrid mice from BXD and Tagl-Cre crosses, chosen to represent a broad range of neuronal and glial cell densities (Lu et al., 2001; Mozhui et al., 2007; Seecharan et al., 2003). Each brain first underwent multi-shell *ex vivo* dMRI (150µm isotropic resolution), enabling estimation of tissue microstructure using both phenomenological models, such as Diffusion Kurtosis Imaging (DKI, Jensen et al., 2005), and multi-compartment biophysical models, such as Soma And Neurite Density Imaging (SANDI, Palombo et al., 2020). Subsequently, the same brains were optically cleared, had dual immunolabelling for neurons and astrocytes and were imaged using whole-brain 3D LSM at cellular resolution (1x1x4µm). Co-registration of the two modalities (Huszar et al., 2023) provided spatially matched MRI and 3D microscopy throughout each brain, forming the basis for testing relationships between dMRI contrasts and underlying cellular features across mouse strains.

A key requirement of this workflow was to make whole-brain tissue processing sufficiently rapid and reproducible for systematic integration with MRI. We therefore implemented *active* tissue processing using an electrophoretic device (Smartbatch+, LifeCanvas Technologies) and compatible clearing and immunolabelling protocols (Kim et al., 2015; Park et al., 2019; Yun et al., 2025). The device applies a rotating electric field to rapidly disperse charged molecules, such as antibodies, throughout the tissue in a stochastic diffusion-like manner (Fig. 1B). This significantly accelerates individual processing steps, particularly immunolabelling, while promoting uniform treatment throughout large tissue volumes. We found that this approach (Yun et al., 2025) was applicable not only to mouse brains but also to substantially larger rat brains (Fig. 1C), extending the practical scale of active tissue processing for whole-brain mapping. Importantly, the processed samples showed no evidence of anisotropic tissue swelling, which has previously been associated with sodium dodecyl sulphate-based delipidation (Johnson et al., 2023).

We further established that MRI contrast enhancement was compatible with subsequent tissue clearing and microscopy. Brains perfused with a gadolinium-based MRI contrast agent retained optical transparency after clearing (Fig. 1D), enabling light penetration throughout the whole-brain volume (Fig. 1E). Gadolinium-enhanced perfusion was also compatible with subsequent immunolabelling (Fig. 2B-D). The workflow can therefore incorporate contrast-enhanced *ex vivo* MRI, which reduces tissue T_1_ relaxation times and allows for shorter MRI acquisitions (D’Arceuil et al., 2007; Kim et al., 2009).

**FIGURE 2.**
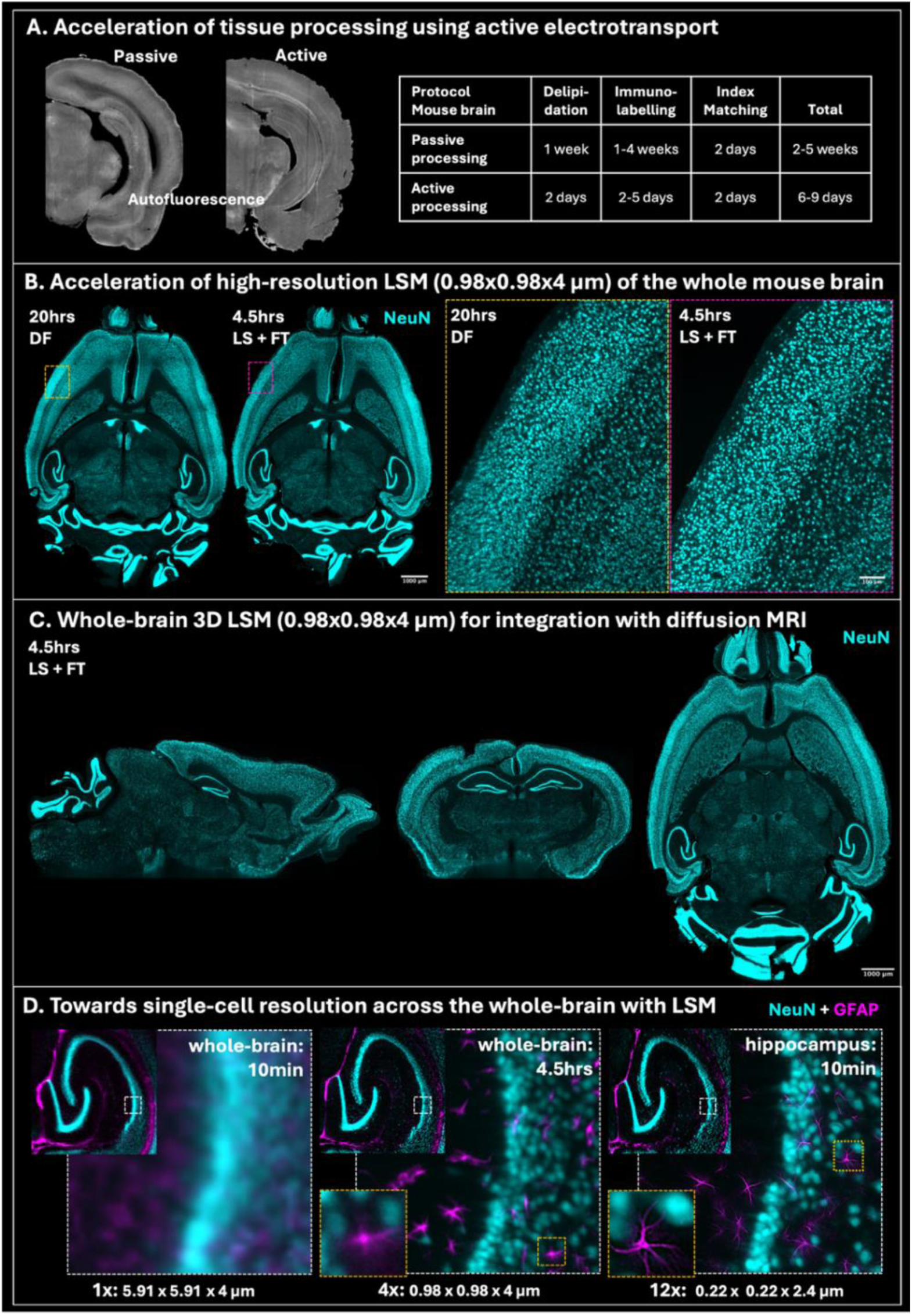
Accelerated tissue processing and light-sheet microscopy (LSM) enable whole-brain imaging at higher throughput. **(A)** Left: Representative coronal views of rat brain hemispheres following tissue processing by passive diffusion for 40 days or active electrotransport for 16 days, showing comparable autofluorescence. Right: Reduction in tissue processing time achieved by active electrotransport compared with passive diffusion in the mouse brain. **(B)** Comparison of Light-speed and fast-tiling (LS+FT) acquisition with conventional dynamic focus (DF) imaging (11 steps) of the same mouse brain using matched acquisition settings (cyan: NeuN-labelled neuronal nuclei). Enlarged regions demonstrate reduced image blurring with the LS+FT acquisition. **(C)** Tissue clearing combined with LSM enables comprehensive 3D imaging of intact rodent brains and facilitates direct integration with MRI. Immunolabelling was successfully achieved following perfusion with gadolinium contrast agent. **(D)** Enlarged views of the right hippocampus acquired using 1x, 4x, and 12x objectives, with approximate acquisition times for whole-brain and hippocampal imaging indicated (magenta: GFAP-labelled astrocytes). High-resolution imaging of a 2mm^3^ region of the hippocampus can be completed in 10minutes using a 12x objective. At a resolution of 0.22x0.22x2.4µm, fine cellular details, including astrocyte processes, are clearly resolved. Deconvolution was performed on images acquired using the 12x objective.

Together, these results establish the key components of the proposed integrative workflow in which the same intact brain can be interrogated consecutively from MRI-scale to cellular-scale resolution. We next optimised each stage of the workflow to increase throughput, enable multiplexed whole-brain cellular mapping, and quantitatively integrate microscopy-derived cellular features with dMRI contrasts across spatial scales.

### Accelerated tissue processing and whole-brain 3D LSM enable high-throughput cellular mapping at micrometre resolution

We addressed two major bottlenecks limiting the scalability of whole-brain 3D microscopy: the time required for uniform tissue processing and the time required for volumetric LSM acquisition. Replacing passive molecular diffusion with active stochastic electrotransport for delipidation and immunolabelling substantially reduced processing times while maintaining effective tissue clearing and whole-brain labelling (Fig. 2A). The mouse-brain protocol, including tissue preservation, delipidation, bleaching, immunolabelling, dehydration, refractive index matching, and all intermediary washes, was completed in under two weeks. The largest reduction arose from active immunolabelling, which required only 2-5 days across the antibodies tested, depending on whether sequential primary and secondary labelling steps were required. By comparison, passive whole-brain immunolabelling typically requires 1-4 weeks and may still result in incomplete or non-uniform labelling (Weiss et al., 2021).

We next accelerated whole-brain image acquisition, by optimising acquisition protocols on a commercial light sheet microscope (Ultramicroscope BLAZE, Miltenyi Biotec). Using LightSpeed (LS) mode, in which stage motion is continuous and synchronised with image acquisition (Price and Lyman, 2024), we acquired a three-channel whole-mouse-brain scan at moderate resolution (5.91x5.91x4µm) in under 10 minutes (Supplementary Fig. 1A). This represents a 30-fold reduction in acquisition time compared with conventional dynamic focus (DF) imaging, where the stage moves incrementally between each step, and which required approximately 3 hours for the same scan. The resulting ultrafast whole-brain acquisition provides a rapid way of assessing tissue processing and labelling quality, before higher resolution microscopy.

For cellular-resolution imaging (0.98x0.98x4µm), we combined LightSpeed mode with fast-tiling (LS+FT) and 3x scaling, using selective exposures to minimise collection of redundant data (Bigott et al., 2026). This enabled acquisition of a large-scale two-channel whole-brain dataset (∼1 TB) in under 5hrs (Fig. 2B, NeuN staining shown as an example), compared with approximately 20 hours using conventional DF imaging. Fast-tiling additionally reduced image blurring by keeping the camera’s field of view fixed relative to the light-sheet focus during sample translation.

The combination of accelerated tissue processing and whole-brain imaging substantially increased the throughput of 3D cellular mapping. Using this workflow, brains from all eight strains were processed within six weeks, with whole-brain LSM acquisition times ranging from under 10min at moderate resolution (5.91x5.91x4µm) to approximately 5 hours at cellular resolution (0.98x0.98x4µm). These timescales approach those of conventional histology and 2D microscopy, while retaining non-destructive 3D imaging of the intact brain (Fig. 2C). They enable measurements across anatomical axes and brain regions, without the constraints imposed by tissue sectioning and sparse sampling inherent to histology.

Importantly, the workflow can also be extended to sub-micrometre resolution (0.22x0.22x2.45µm) across the whole brain, given sufficient acquisition time (e.g., a weekend) and storage capacity. To demonstrate the additional information accessible at this scale, we imaged the right hippocampus at this resolution (Fig. 2D), revealing detailed cellular morphology while preserving its 3D anatomical context. Notably, comparable image quality was also obtained across different light-sheet microscopes and imaging sites (Supplementary Fig. 1B), demonstrating reproducibility across imaging systems. In summary, the proposed workflow provides a flexible acquisition strategy spanning rapid whole-brain mesoscale screening, to detailed cellular-resolution mapping.

### Whole-brain multiplexed immunolabelling enables simultaneous mapping of multiple cell types

We next assessed whether our active immunolabelling setup could provide specific labelling of different cellular populations throughout the intact mouse brain (Fig. 3A; Supplementary Fig. 2 and 3). Using the same tissue processing framework, we achieved whole-brain labelling of neurons (NeuN - neuronal nuclei), astrocytes (GFAP - glial fibrillary acidic protein), oligodendrocytes (MOG - myelin oligodendrocyte glycoprotein), microglia (IBA-1 - ionized calcium-binding adaptor molecule 1), and dopaminergic neurons (TH - tyrosine hydroxylase), with active staining completed within 5 days. Labelling was achieved using commercially available conventional antibodies, rather than using specialised probes, making the approach readily adaptable to other tissues and targets. Labelled cells were detected throughout the depth of the brain with low levels of background signal (Fig. 3, Suppl. Fig. 2, 3). Moreover, we observed no obvious staining gradients or surface-restricted labelling, features commonly associated with passive immunolabelling approaches (Kim et al., 2015).

**FIGURE 3.**
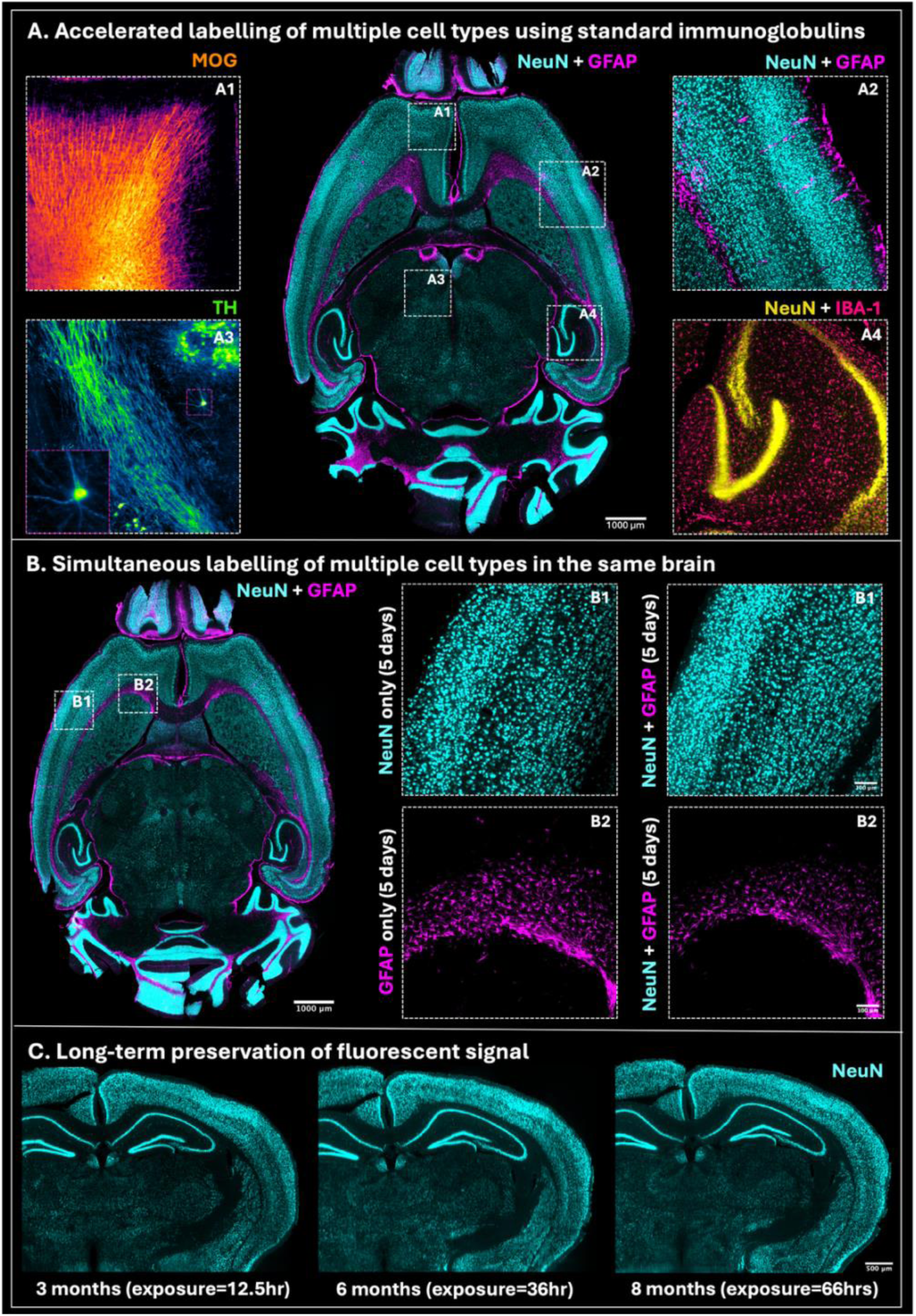
Whole-brain multiplexed immunolabelling enables mapping of different cell types in intact mouse brains. **(A)** Centre: Axial view of a gadolinium-perfused mouse brain following dual immunolabelling of neurons and astrocytes. Left and right: Enlarged views of representative regions from separate brains labelled with the indicated markers; approximate locations are shown on the axial image. **(A1)** MOG, labelling oligodendrocytes and myelin. **(A2)** NeuN and GFAP dual labelling of neuronal cell bodies and astrocytes. **(A3)** TH, labelling dopaminergic neurons and their projections. **(A4)** NeuN and IBA-1 dual labelling of neuronal nuclei and microglia. **(B)** Left: Axial view of mouse brain dual-labelled for neurons and astrocytes. Right: Enlarged views of representative regions from brains subjected to single- or dual-marker immunolabelling, with approximate locations shown on the axial image, demonstrating that simultaneous labelling of neurons and astrocytes did not increase processing time or non-specific background signal. **(C)** Partial coronal view of the same mouse brain imaged over six sessions spanning several months, with three representative sessions distributed across the imaging period shown alongside the cumulative imaging times. Fluorescence signal was preserved despite repeated and prolonged illumination. Samples were stored at room temperature in ethyl cinnamate.

By pairing each cellular target with a spectrally distinct fluorescent antibody, we achieved multiplexed active labelling of multiple cell types in the same brain. Dual immunolabelling of neurons (NeuN) and astrocytes (GFAP) simultaneously within the same sample yielded contrasts comparable to those obtained from single-target labelling in separate specimens (Fig. 3B). Because both targets were labelled concurrently, multiplexing did not increase processing time. Further expansion of the number of simultaneously labelled targets is possible, although multiplexing capacity is currently limited by tissue autofluorescence, light absorption and scattering at shorter wavelengths; reduced detector sensitivity at longer wavelengths; and the availability of compatible laser lines and filter sets.

The protocol was also compatible with long-term sample storage. Successful NeuN, GFAP and TH immunolabelling was obtained in mouse brains >12 months after perfusion-fixation (Fig. 3A and Supplementary Figs. 1 and 2). The fluorescence signals remained stable for more than 6 months after labelling, and were preserved despite prolonged and repeated illumination, enabling further microscopy analyses of the same labelled samples (Fig. 3C). Together, these developments provide a versatile platform for whole-brain cellular phenotyping at scale.

### Combination of diffusion MRI contrasts is predictive of LSM-derived cellular features across genetically diverse mice

To directly relate dMRI-derived microstructural measures to cellular features, we integrated quantitative whole-brain LSM measurements and dMRI contrasts within the same spatial framework (Fig. 4). Whole-brain, high resolution imaging of dual-labelled brains (0.98x0.98x4µm) generated approximately 1TB of LSM data per specimen. Neuronal nuclei were segmented from NeuN-labelled images to generate whole-brain maps of neuronal density (Fig. 4A, Supplementary Fig. 4). GFAP-positive stain area fraction (SAF) was quantified as a proxy for astrocytic content, because individual astrocytes could not be reliably delineated at this acquisition resolution. In parallel, multi-shell dMRI data (150µm isotropic resolution, six shells up to b=17.5 ms/µm^2^) were processed to derive phenomenological DKI measures (Jensen et al., 2005) and biophysical parameters from the SANDI model (Palombo et al., 2020), including neurite and soma volume fractions (f_neurite_ and f_soma_ respectively).

**FIGURE 4.**
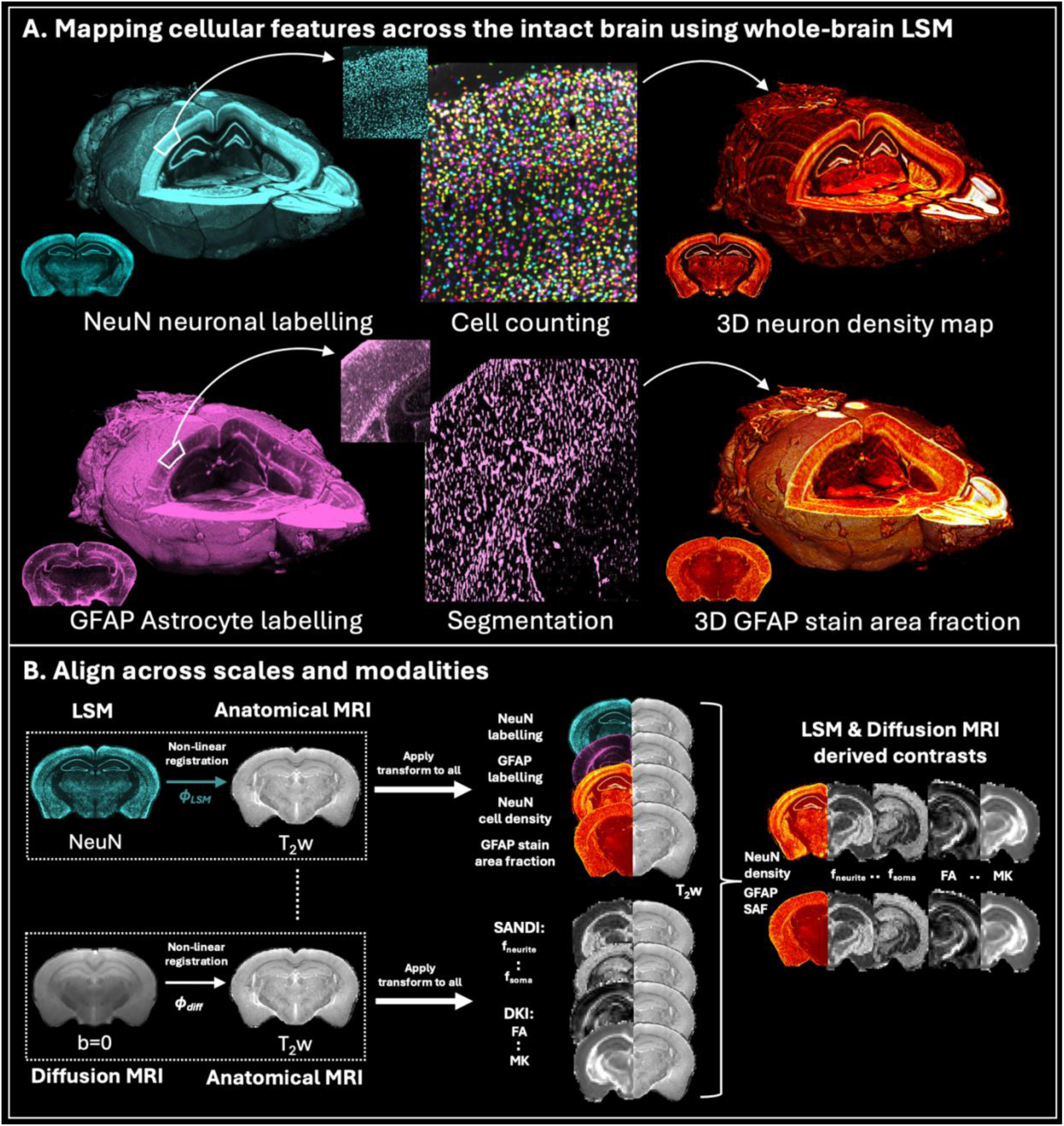
Extraction of cellular features from 3D light-sheet microscopy data and integration with MRI. **(A)** Top: Segmentation of neuronal cell density and astrocytic content from LSM data. Neuronal nuclei were segmented using Cellpose from the NeuN labelled channel. The segmented cells were counted to generate a 3D map of neuronal density across the whole-brain. Bottom: Astrocytic context was obtained from the GFAP labelled channel. GFAP-positive pixels were segmented using Ilastik. The fraction of tissue occupied by GFAP-positive signal (GFAP stain area fraction - SAF) was used as a proxy of astrocytic content. **(B)** Microscopy and MRI data acquired from the same mouse were anatomically aligned to enable spatially-matched comparison across modalities. NeuN-labelled LSM data for each sample was non-linear registered to the corresponding T_2_w MRI. The calculated transformation (φ_LSM_) was applied to the NeuN- and GFAP-labelled LSM data, as well as the neuron density and GFAP SAF, to align these contrasts with the corresponding T_2_w MRI. The diffusion MRI data were independently aligned to the T_2_w MRI using a separate nonlinear registration (φ_diff_). This process generated a multi-modal volumetric dataset for each mouse using the corresponding T2w MRI as common reference, enabling direct voxel- and region-wise comparisons across imaging modalities and spatial scales.

For each of the eight brains, the resulting microscopy and dMRI maps were non-linearly co-registered in 3D, using the corresponding anatomical T2-weighted MRI as a common reference (Fig. 4B; Supplementary Figs. 5 and 6). This generated spatially matched multimodal datasets at the dMRI resolution (150 µm isotropic), enabling direct voxel-wise and region-wise comparisons between MRI-derived microstructural measures and LSM-derived neuronal density and GFAP SAF, within the same brain and across the eight animals.

We used these spatially matched datasets to assess whether individual dMRI contrasts or combinations of them could predict cellular features measured directly by LSM (Fig. 5). For multivariate prediction, voxel-wise piecewise linear random forests (RF) models were trained independently for each animal (each from a separate mouse strain), using 80% of voxels for training and evaluated on the remaining 20%. LSM-derived NeuN neuronal density and GFAP SAF were used as dependent variables, with SANDI and DKI measures as regressors (Fig. 5A).

**FIGURE 5.**
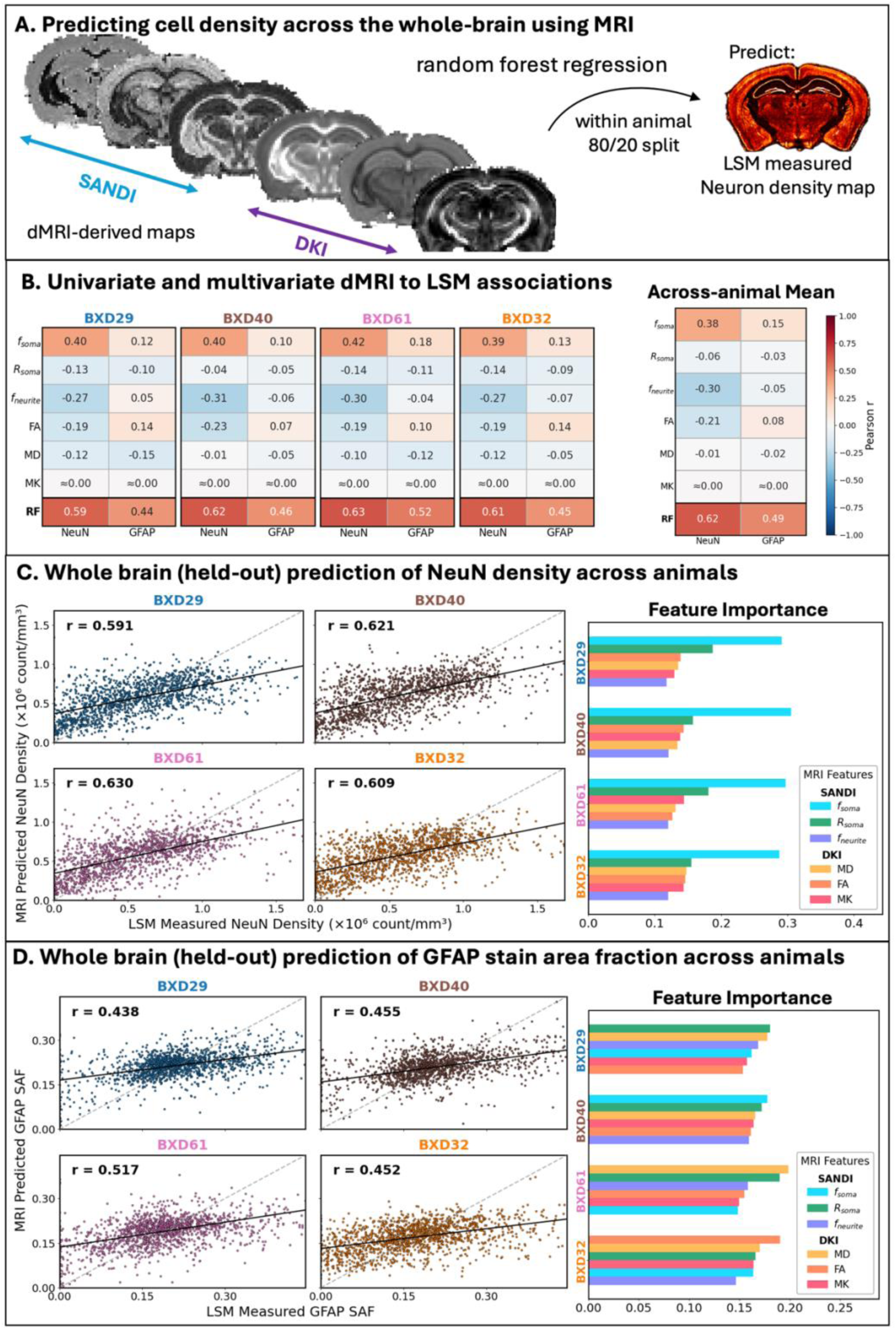
Predicting LSM-derived neuronal cell density and astrocyte stain area fraction (SAF) using MRI. **(A)** Following alignment of LSM and dMRI, random forest models were trained to test whether LSM-derived neuronal density maps or astrocyte SAF could be accurately predicted from dMRI-derived metrics (SANDI and DKI). The random forest models were trained and tested using a voxel-wise, within brain and within animal 80/20 train/test split. **(B)** Correlation of individual dMRI SANDI and DKI contrasts against LSM-derived NeuN neuronal density and GFAP SAF for four animals (each from a different strain). Bottom row shows the regression coefficient of the multivariate random forest (RF) model that considers all dMRI metrics against the same LSM-derived cellular features. The mean correlations across all eight animals are presented on the right. Supplementary Fig. 7 shows associations for all eight animals. **(C, D)** Predictions of neuronal cell density and GFAP SAF respectively, evaluated on held out data, across four animals, representing the worst, best and two intermediate R scores. Relative feature importance for SANDI and DKI metrics is shown for each strain. Supplementary Fig. 8 shows performance for all animals.

Individual dMRI measures generally showed weak-to-moderate correlations with LSM-derived cellular maps (Fig. 5B, Supplementary Fig. 7A). Notably, the SANDI parameter f_soma_ was the individual measure with the strongest association with NeuN neuronal density (mean Pearson’s r= 0.38 across the eight mice). Combining dMRI measures substantially improved predictive performance. Voxel-wise neuronal density maps predicted using all SANDI and DKI measures more closely reproduced the LSM-derived maps than predictions based on f_soma_ alone (Supplementary Fig. 7B). Across all animals, the multivariate RF models explained on average 35% of variance in NeuN neuronal density for held-out voxels (mean R≈0.6, Fig. 5C). Multiple dMRI measures contributed to prediction, with f_soma_ consistently showing the highest feature importance across animals (Supplementary Fig. 8A). Phenomenological DKI measures, including fractional anisotropy (FA), mean diffusivity (MD) and mean kurtosis (MK), also contributed to model performance, suggesting that information relevant to neuronal density is distributed across multiple dMRI contrasts and is not fully captured by a single contrast or biophysical parameter.

Prediction of GFAP SAF was weaker than that of neuronal density (R ≈ 0.5), and the relative importance of dMRI microstructural features was more heterogeneous across animals (Fig. 5D, Supplementary Fig. 8B). The greater and more consistent contribution of SANDI f_soma_, to neuronal-density prediction than to GFAP SAF suggests differential sensitivity of individual dMRI parameters to distinct cellular features. Nevertheless, combining dMRI measures also improved prediction of astrocytic content relative to individual measures (Supplementary Fig. 7A), demonstrating that complementary information across diffusion contrasts extends beyond neuronal density.

We additionally assessed the extent to which the MRI–LSM relationship transferred across animals (Supplementary Fig. 9). First, mean neuronal-density predictions from held-out voxels of the independently trained within-animal models were compared with the corresponding mean LSM-derived neuronal densities across the eight brains (Supplementary Fig. 9A). Across gray matter, mean MRI-based predictions closely tracked the corresponding mean LSM-derived measurements. We then performed a more stringent leave-one-animal-out analysis, in which a multivariate RF model was trained using pooled voxel-wise MRI and microscopy measurements from seven brains and applied to the eighth brain, which was entirely excluded from model training. This was repeated eight times, so that each animal served once as the held-out test case. Mean grey-matter neuronal-density predictions in the held-out animals showed an association with the corresponding LSM-derived measurements (R=0.63, Supplementary Fig. 9B). Although between-animal prediction was weaker than within-animal prediction, the multivariate dMRI– cellular relationship retained predictive information when transferred to previously unseen brains.

Taken together, these results demonstrate that complementary information distributed across multiple dMRI contrasts (SANDI and DKI in this study) can be integrated to improve prediction of cellular features, beyond the prediction performance achieved by individual contrasts. More broadly, our proposed cross-scale framework provides a means of identifying and testing microstructural MRI measures and their combinations that more closely reflect underlying cellular architecture.

## Discussion

We have established a platform for cross-scale imaging of the same intact rodent brain using 3D light-sheet microscopy (LSM) and diffusion MRI (dMRI). By combining accelerated tissue clearing, active immunolabelling, and high-throughput volumetric microscopy we achieved whole-brain cellular mapping on timescales approaching conventional histology, while preserving the 3D organisation of the intact brain. This enables direct spatial integration of cellular measurements with MRI-derived microstructural contrasts acquired from the same specimen. Whole-brain immunolabelling further enables simultaneous visualisation of multiple targets within the same specimen, allowing for a diverse range of cellular and microscopical features to be investigated in a common anatomical framework. Applying this integrative approach across genetically diverse mice, we have shown that combinations of dMRI contrasts predict neuronal and astrocytic features more accurately than individual contrasts. A complementary cross-animal analysis further indicates that part of the dMRI–cellular relationship is retained when prediction is extended to previously unseen brains. Our findings establish the proposed platform as a scalable *ex vivo* testbed for evaluating, calibrating and combining dMRI contrasts against cellular measurements, with the goal of guiding the development of new MRI measures that can be ultimately translated to living brains, including human ones.

A major challenge for whole-brain microscopy is balancing tissue coverage, molecular specificity, and acquisition throughput. Here, we addressed these limitations through electrotransport-based tissue processing, active whole-brain immunolabelling and accelerated LSM acquisition strategies. Using conventional antibodies and a standardised protocol, we demonstrated robust and generalisable whole-brain immunolabelling across multiple cellular targets, including neurons, astrocytes, oligodendrocytes, microglia, and dopaminergic neurons. Unlike approaches relying on genetically encoded fluorescent reporters (e.g., Thy1-eYFP) or viral tracers (Gimenez et al., 2017; Goubran et al., 2019; Stolp et al., 2018), antibody-based labelling decouples signal generation from genetically encoded or virus-mediated labels. This expands the range of cellular and molecular targets that can be interrogated and, importantly, enables molecular phenotyping in tissues for which genetic labelling strategies are unavailable, supporting translation to human tissue (Park et al., 2024). Antibody penetration and labelling were achieved throughout the intact brain with low background signal and without the staining gradients commonly associated with passive diffusion. Together, this demonstrates the feasibility of simultaneous, scalable, and broadly generalisable molecular phenotyping across the whole brain using antibody-based methods. Combined with batch processing during active immunolabelling (the SmartBatch+ can accommodate up to 12 mouse or 4 rat brains per run), these features make systematic MRI-microscopy studies across multiple specimens increasingly practical.

A key strength of this study is the integration of multimodal imaging across micro- and meso-scopic scales (0.98-150µm). By co-registering high-resolution 3D LSM data with MRI acquired from the same brain, we enabled direct voxel- and region-wise comparisons between cellular organisation and diffusion MRI-derived measures of tissue microstructure. Individual dMRI measures showed only weak-to-moderate relationships with microscopy-derived cellular features, despite some measures being designed to capture specific aspects of tissue microstructure. Among the measures examined here, the SANDI soma volume fraction, f_soma_, showed the strongest individual association with neuronal density (Pearson’s r∼0.38) and was consistently the most informative feature in multivariate models of neuronal density. Nevertheless, combining f_soma_ with complementary SANDI and DKI measures improved prediction substantially, with multivariate models achieving mean held-out correlations of approximately R∼0.6. These findings motivate the development of MRI protocols and models that optimise combinations of complementary measurements rather than relying exclusively on individual parameters that aim for one-to-one mapping with underlying cellular features.

The difference between neuronal and astrocytic predictions further illustrates the complexity of MRI–cellular relationships. Prediction of GFAP stain area fraction (SAF) was weaker than prediction of neuronal density, and the relative contributions of individual dMRI measures were more heterogeneous across strains. In particular, f_soma_ contributed more consistently to prediction of neuronal density than GFAP SAF, suggesting differential sensitivity of individual dMRI parameters to distinct cellular features. This demonstrates the value of multiplexed cellular measurements our LSM setup provides for determining which aspects of tissue composition contribute to MRI contrasts.

The associations identified between dMRI and microscopy measurements using within-animal models preserved differences in mean neuronal density across the eight genetically distinct animals. A between-animal leave-one-out analysis provided a more stringent assessment of the generalisability of these relationships. Performance was lower than for models trained and tested within the same brain, potentially reflecting both inter-animal and inter-scan variability factors (such as temperature differences or specimen condition). Nevertheless, the persistence of a relationship in completely held-out animals suggests that at least part of it is shared across brains. As each genetic strain was represented by a single animal, the current data do not permit inference about strain-specific or genetic effects and replication with larger cohorts will be required.

The ability to integrate MRI with whole-brain cellular mapping as demonstrated here can have broader implications for neuroscience research. Whole-brain approaches are particularly valuable for studying processes that involve distributed networks rather than isolated anatomical regions, including neurodevelopment, neurodegeneration, neuroinflammation, and brain-wide connectivity changes. By preserving intact tissue architecture and mesoscale connectivity while simultaneously enabling cellular-resolution imaging, this framework provides a powerful platform for multiscale brain mapping.

Previous studies have demonstrated efforts in linking diffusion MRI with LSM by relying on data acquired from different animals and registration to anatomical atlases (Goubran et al., 2019; Han et al., 2026), limiting direct correspondence between cellular and MRI measurements within the same brain. Other studies have demonstrated the potential of cross-scale imaging within the same tissue samples (Leuze et al., 2021; Stolp et al., 2018), but have not extended this approach to intact whole brains. More recently, whole-brain approaches (Johnson et al., 2023; Mansour et al., 2025) have integrated high-resolution MRI and LSM within the same specimens, enabling multimodal mapping and construction of stereotaxic cellular atlases. In these studies, high-resolution MRI provided a geometrically faithful reference for LSM data affected by tissue-processing deformations. Our approach addresses a complementary challenge by making quantitative whole-brain cellular microscopy sufficiently rapid and scalable, to overcome tissue processing and microscopy bottlenecks, and serve as a biological reference for interpreting microstructural dMRI of the same brain. To our knowledge, this is the first application of whole-brain LSM to directly investigate the relationships between dMRI-derived microstructural measures and cellular features of both neurons and astrocytes, with potential of expanding further (de Guzman et al., 2026). By combining active tissue processing, multiplexed immunolabelling, accelerated whole-brain LSM, and spatially matched multi-shell dMRI, we directly test how microscopy-derived cellular features of these two major cell classes are represented in dMRI contrasts. This approach extends beyond multimodal anatomical integration, towards evaluating and refining the cellular specificity of microstructural MRI models, including SANDI.

Despite the strengths of our approach, several limitations should be considered. Firstly, the predictive models for each strain were trained and evaluated using randomly-selected voxels within individual brains. Even if held-out voxels were not used for model fitting, this approach does not provide a fully independent test dataset. Secondly, the single animal per strain limits explorations on capturing inter-strain variability. Nevertheless, the throughput established here provides a route to addressing these limitations in larger studies. Thirdly, translation from *ex vivo* MRI measurements to *in vivo* MRI presents additional challenges. Fixation and the postmortem state inevitably alter several properties relevant to diffusion measurements, including diffusion coefficients of different tissue compartments, cell membrane permeability, compartment volume fractions and tissue size. The use of gadolinium-based contrast agents during perfusion fixation represents a further consideration, as gadolinium alters relaxation properties of tissue and therefore MRI-derived measurements. Even if *ex vivo* measurements do not provide a direct quantitative representation of *in vivo* tissue properties, they allow the relationship between MRI measurements, biophysical models and underlying tissue microstructure to be systematically explored. Our setup allows *in vivo* MRI scans to be incorporated into the analysis, and in fact we are already working towards this direction (Wood et al., 2026).

A related question concerns the translation of findings from rodents to humans. Rodent models provide several important advantages for the development and validation of advanced MRI methods. Tissue processing protocols are well established and standardised. *Ex vivo* imaging of their small brains with high-field small bore scanners and cryogenic probes produces high-resolution data with high signal-to-noise ratio, while perfusion fixation allows excellent preservation of tissue structure and facilitates direct comparison between MRI and histological measurements. In addition, several aspects of brain tissue microstructure are preserved across mammalian species. These include, for instance, the dense packing of neuronal and glial processes, extracellular-space volume fraction and non-neuronal cell density (Herculano-Houzel et al., 2007, 2006; Syková and Nicholson, 2008). Although absolute neuronal density and astrocytic domain size differ substantially between rodents and humans, many MRI contrasts are determined less by cell number and more by the aggregate volume fractions, molecular environments, and spatial organization of cellular and extracellular compartments (Lampinen et al., 2023). Consequently, microscopy validation of MRI in rodents can establish the biological sensitivity and mechanistic interpretation of an MRI contrast when applied to humans, even where individual cellular dimensions differ between species.

In summary, we have established a scalable cross-resolution framework that links whole-brain 3D light-sheet microscopy directly with diffusion MRI in the same specimens. The combination of accelerated tissue processing, multiplexed immunolabelling, rapid volumetric microscopy, and multimodal registration enables cellular architecture to be quantified across intact brains and related directly to MRI-derived microstructural measures. Our findings show that information on cellular features is distributed across multiple dMRI contrasts and that combining complementary measures improves prediction of neuronal and astrocytic features across eight genetically distinct animals. More broadly, this framework provides a platform for using quantitative 3D microscopy to test, refine, and optimise microstructural MRI measures, with the longer-term goal of developing more biologically-specific imaging markers that can be applied non-invasively in the living human brain.

## Methods

### Animals

Eight adult male mice from genetically distinct F1 hybrid mice from BXD and Tagl-Cre crosses (Table 1) underwent *ex vivo* MRI followed by 3D LSM. All procedures were conducted in accordance with the UK Animals (Scientific Procedures) Act 1986 and approved by the University of Oxford’s ACER (Animal Care and Ethical Review) committee (22 October 2024).

**TABLE 1.** Mouse strains and sex.

| **Strain** | **Sex (count)** |
|---|---|
| BXD12/TyJ; Tagln-cre | M (1) |
| BXD29/TyJ; Tagln-cre | M (1) |
| BXD32/TyJ; Tagln-cre | M (1) |
| BXD34/TyJ; Tagln-cre | M (1) |
| BXD40/TyJ; Tagln-cre | M (1) |
| BXD61/RwwJ; Tagln-cre | M (1) |
| BXD89/RwwJ; Tagln-cre | M (1) |
| B6.Cg-Tg(Tagln-cre)1Her/J (Lerch, J) | M (1) |

All animals were anaesthetised with pentobarbital (Dolethal) and then transcardially perfused with phosphate-buffered saline (PBS), followed by 4% paraformaldehyde (PFA) in PBS according to an established protocol (Cahill et al., 2012; de Guzman et al., 2016; Smart et al., 2023). Brains were retained within the skull and post-fixed overnight in 4% PFA at 4°C before being transferred to 0.01M PBS (Invitrogen, AM9625) containing 0.02% sodium azide (Sigma-Aldrich, S8032) for storage at 4°C. For all mice, 2mM gadolinium-based contrast agent (Gadovist, Bayer) was added to the perfusate, post-fixative and storage solution. Gadolinium, shortens the longitudinal relaxation time (T_1_), thereby accelerating signal recovery and enabling the use of shorter repetition times (TR) during MRI.

High resolution MRI was performed two weeks after perfusion. Following MRI, brains were extracted, immunolabelled, and rendered optically transparent for whole-brain LSM. An overview of the experimental workflow is shown in Fig. 1A.

To explore generalisability and optimise tissue clearing protocols, two adult male Sprague Dawley rats (Charles River, Margate, UK) also underwent a similar procedure, with transcardial perfusion, fixation and storage. For one of the rats, 2mM gadolinium-based contrast agent was added to the perfusate, post-fixative and storage solution, while no gadolinium contrast agent was used for the second.

### MRI

All mice underwent *ex vivo* MRI on a 7T Bruker BioSpec 70/20 USR scanner with a four-channel cryocoil and 660mT/m gradients. Samples were kept at room temperature (18-22°C) throughout the acquisition. The acquisition protocol included high-resolution T_2_-weighted structural imaging (60µm isotropic) and multi-shell diffusion MRI (150µm isotropic). T_2_-weighted images were acquired using a 3D TurboRARE sequence (TE/TR= 36/350ms, RARE factor=6, ESP=12ms, BW=60 kHz). Multi-shell dMRI data were acquired using a pulsed-gradient spin-echo sequence with a 12-segment 3D-EPI readout, 16 directions per shell (b = [1, 2.5, 5.5, 8.5, 12.5, 17.5] ms/µm^2^, *δ*/Δ=7.5/20ms), and interleaved b=0 images. The total acquisition time was approximately 12 hours per animal.

### Tissue clearing and immunolabelling

Active tissue clearing and immunolabelling were performed using previously described methods (Yun et al., 2025) with a SmartBatch+ device (LifeCanvas Technologies). Unless stated otherwise, reagent volumes of 20ml and 40ml were used per mouse or rat brain, respectively. Washes were performed in 40ml of 0.01M PBS containing 0.02% sodium azide.

Following MRI, whole brains were removed from the skull and preserved using SHIELD (LifeCanvas Technologies; Park et al., 2019). Mouse brains were incubated in SHIELD OFF reagent at 4°C for 3 days, followed by SHIELD ON reagent at 37°C for 24hrs with mild agitation. For rat brains, the SHIELD OFF incubation was extended to 6 days.

Mouse brains were preincubated in Clear+ Delipidation Buffer (LifeCanvas Technologies) at 45°C for at least 24hrs before active delipidation in the SmartBatch+ device for 30-40hrs. For rat brains, active delipidation was extended to 6 days. Brains were subsequently bleached overnight at 4°C in 5% hydrogen peroxide (Fisher H/1800/15).

For immunolabelling, mouse brains were preincubated in Primary Sample Buffer (LifeCanvas Technologies) overnight at room temperature, followed by an additional incubation of at least 3hrs. Active primary antibody labelling was then performed for 18hrs in the SmartBatch+ device (Yun et al., 2025), followed by two 4hr washes and overnight fixation in 4% PFA. Brains were subsequently preincubated in Secondary Sample Buffer (LifeCanvas Technologies) at 37°C for 6-8hrs, with the buffer refreshed halfway through, before active secondary antibody labelling for 12hrs in the SmartBatch+. Following secondary labelling, brains underwent two 3hr washes in Secondary Sample Buffer in the SmartBatch+, followed by two 1hr washes, overnight fixation in 4% PFA, and two final 1hr washes. NeuN and GFAP co-labelling was performed by sequential delivery of the corresponding secondary antibodies (Table 2). The same protocol was used for MOG, and IBA-1. For TH labelling, the primary antibody and Fab fragment were delivered simultaneously, and the secondary labelling steps were omitted.

**TABLE 2.**
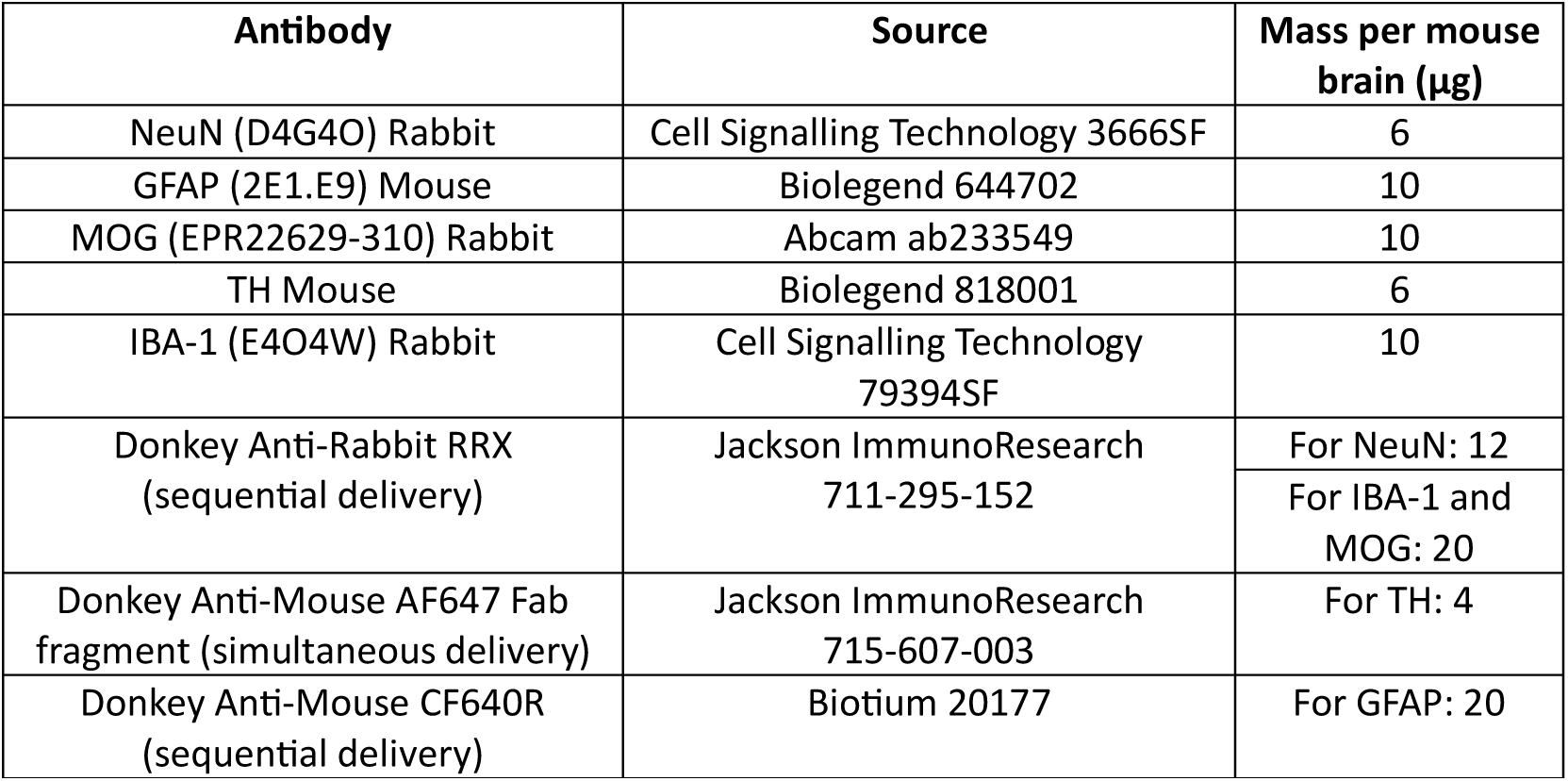
Antibodies used for active immunolabelling.

Following immunolabelling, brains were dehydrated through a graded methanol series (VWR, 20847.307; 20%, 40%, 60%, 80%, and 100%) for 1hr at each concentration, followed by overnight incubation in 100% methanol. Brains were then transferred to ethyl cinnamate (Sigma, 112372-100G) and incubated overnight for refractive index matching (RI=1.56).

### Light-sheet microscopy

Following optically clearing, brains were mounted horizontally using applying superglue applied to the inferior surface and imaged in ethyl cinnamate. Imaging was performed using an Ultramicroscope BLAZE (Miltenyi Biotech), operated using Imspector Pro 7.8.1, and equipped with the Multi-Immersion Planar (MI Plan) objective lens series, which provides magnifications of up to 30x. Lightspeed mode was evaluated against conventional imaging using dynamic horizontal focus (Price and Lyman, 2024).

For whole-brain imaging, the sample was illuminated from both sides (left/right) using three sheets at maximum width (NA=0.163) and minimum thickness (3.9µm). Sample autofluorescence (Ex/Em: 488/520-50nm) and immunolabelling (NeuN Ex/Em: 561/595-40nm; GFAP Ex/Em: 640/680-30nm) were first imaged using the 1.1x objective with 1x zoom. These acquisitions generated moderate-resolution whole-brain image volumes (5.91x5.91x4µm³) for quality control (Suppl. Fig. 1A). Higher-resolution whole-brain volumes (0.98x0.98x4µm³) of immunolabelling were acquired using the 4x objective with 1.66x zoom for subsequent cell segmentation (Figs. 2B, 4A). To visualise cellular morphology, immunolabelling within a restricted region of the right hippocampus (2mm^3^) was imaged at maximum resolution (0.22x0.22x2.45µm^3^) using the 12x objective with 2.5x zoom (Fig. 2D). These high-resolution acquisitions were performed in Deconvolution mode using right-sided illumination with a single light sheet at 30% width (NA=0.60) and a thickness of 6.1µm.

To assess the generalisability of the methodology across microscopy systems and facilities, a single mouse brain was imaged at moderate-resolution on three light-sheet microscopes with differing hardware (e.g., camera and optics) and software (e.g., file formats). These included an additional Ultramicroscope BLAZE and an Ultramicroscope II (Miltenyi Biotech) at the University of Oxford and at the University of Birmingham respectively (Suppl. Fig. 1B). Whole-brain imaging was performed using dynamic horizontal focus (DF) with dual-sided illumination and three light sheets at maximum width and minimum thickness (3.9µm). All acquisitions were performed using three-channels and were completed in under 3hrs. Acquisition parameters are summarised in Table 3.

**TABLE 3.** Parameters for acquisitions.

| **Site** | **Scope** | **Objective** | **Zoom** | **Sheet NA** | **DF steps** | **Resolution (µm)** |
|---|---|---|---|---|---|---|
| **Nottingham** | BLAZE | 1.1 | 1 | 0.163 | 23 | 5.91x5.91x4 |
| **Oxford** | BLAZE | 1.1 | 1 | 0.186 | 13 | 5.91x5.91x4 |
| **Birmingham** | UMII | 2 | 0.63 | 0.09 | 25 | 4.79x4.79x4 |

### Image processing

#### MRI processing

Raw dMRI data from the four coil elements were combined using a root-sum-of-squares approach. The resulting data were intensity normalised, by scaling the signal such that mean intensities of interspersed b=0 images acquired throughout the scan were matched, and subsequently denoised in the complex domain using NORDIC (Moeller et al., 2021). A whole-brain mask was generated by skull-stripping the mean b=0 image using bet4animal from the FMRIB Software Library (FSL), with a fractional intensity threshold of 0.075.

Diffusion MRI data was preprocessed using an established pipeline (Tisca et al., 2023). Gibbs-ringing artefacts were removed using MRtrix (Tournier et al., 2019). Susceptibility-induced distortions were then corrected using a pair of blip-reversed AP-PA b=0 images with FSL’s topup (Andersson et al., 2003), followed by correction of eddy current-induced distortions using eddy (Andersson and Sotiropoulos, 2016). Biophysical models were then fitted to the pre-processed data. Diffusion kurtosis imaging (DKI; Jensen et al., 2005) was used to characterise Gaussian and non-Gaussian diffusion and was fitted to the b = [1, 2.5, 5.5, 8.5] ms/µm^2^ shells. In addition to this phenomenological model, the multi-compartment microstructural model SANDI (Palombo et al., 2020) was fitted using the full acquisition (b = [1, 2.5, 5.5, 8.5, 12.5, 17.5] ms/µm^2^) using the SANDI Matlab toolbox (https://github.com/palombom/SANDI-Matlab-Toolbox-Latest-Release). SANDI was fitted with an additional dot-compartment (f_dot_) to model water that is effectively immobile on the diffusion time scale. The intra-soma diffusivity (D_soma_) was fixed at 2.0 µm²/ms, reflecting the reduced diffusivity observed following tissue fixation.

For each mouse, the diffusion b=0 volumes were registered to the corresponding T_2_-weighted anatomical MRI image using image registration methods optimised for the mouse (Chakravarty et al., 2013; Friedel et al., 2014).

#### LSM processing

For acquisitions performed using the 4x and 12x objectives, image volumes were reconstructed by stitching adjacent tiles with a 10% overlap using MACS iQ View 1.2.0 (Miltenyi Biotec). Optimal tile displacements were determined using cross-correlation. The stitching procedure generated a single 3D pyramidal OME-TIFF file for each imaging channel. No tiling was necessary for the lower-resolution acquisition using the 1x objective. The hippocampal acquisition obtained with the 12x objective was deconvolved using MACS iQ View (Miltenyi Biotec).

After stitching, high-resolution whole-brain images acquired using the 4x objective (0.98x0.98x4μm³) were converted from OME-TIFF to HDF5 format to enable segmentation at native resolution using parallelised, patch-based processing on a GPU. Neuronal nuclei were segmented from whole-brain NeuN-labelled LSM data (4x objective) using a pre-trained Cellpose model (Stringer et al., 2021) that was fine-tuned on a manually annotated subset of the dataset, following Pachitariu and Stringer (2022). Individual neuronal nuclei were identified from the resulting segmentation masks, and their centroids were retained (Suppl. Fig. 4). GFAP-positive signal was also segmented in native resolution from whole-brain LSM images (4x objective) using ilastik (Berg et al., 2019). Individual astrocyte segmentation was more challenging than neuronal segmentation at this whole-brain resolution. We therefore quantified the fraction of GFAP-positive foreground pixels within each voxel or region as the stain area fraction (SAF; Kor et al., 2022), which was used as a proxy for astrocytic content (Suppl. Fig. 4). After segmentation, whole-brain neuronal density and GFAP-positive stain area fraction maps were downsampled to a voxel size of 32μm^3^ and then exported in NIfTI format.

To facilitate downstream processing and integration with MRI data (e.g. registration), the whole-brain 4x OME-TIFF volumes were downsampled in Python by block averaging using integer downsampling factors of 32x32x8, resulting in a voxel size of ∼(30μm)^3^. After image reorientation and assignment of appropriate spatial metadata (Left-Right, Inferior-Superior etc), the downsampled volumes were exported to NIfTI format.

#### MRI and LSM registration

The whole-brain, multi-channel LSM images were aligned to the corresponding anatomical T_2_-weighted (T_2_w) MRI in 3D. Although both imaging modalities depict the same intact brain, nonlinear registration was used to account for differences in brain shape and size arising from removal from the skull and subsequent tissue-processing steps. Given that a transformation for aligning diffusion MRI data to the T_2_w images is obtained in the MRI pre-processing pipeline, the LSM to T_2_w transformation also ensures LSM and dMRI alignment.

The downsampled LSM channel of NeuN-staining (30μm^3^) was background thresholded (using the median intensity) and square-rooted. The resulting image gave a similar contrast to anatomical MRI and was used to estimate a non-linear registration warp between the LSM space and the corresponding anatomical T_2_w MRI space of a given sample. This was done using the Tensor Image Registration Library (TIRL, Huszar et al., 2023), designed for microscopy to MRI registration, following a sequence of rigid-fine, anisotropic scaling, affine and nonlinear registration steps (Suppl. Fig. 5, 6). The resulting registration warp enabled alignment for the rest of LSM data and channels (NeuN-staining, GFAP-staining, neuron density and GFAP stain area fraction maps) to the anatomical T_2_w MRI (Suppl. Fig. 5, 6).

For further processing, both LSM and dMRI data were brought to each animal’s T_2_w space and the LSM data resampled to the dMRI resolution (150x150x150 µm^3^) using trilinear interpolation. This enabled voxel-wise comparison of LSM and dMRI measurements within the same sample for the multiple mouse strains, building on previous work integrating MRI with histology (Tisca et al., 2024).

### Modelling across imaging modalities

We tested how accurately the whole-brain distributions of neuronal cell density and astrocyte SAF could be predicted from a given set of dMRI-derived DKI and SANDI maps. Separate random-forest regression models were trained for each strain. For each voxel, the input features comprised FA, MD, and MK from DKI, together with soma volume fraction (f_soma_), soma radius (R_soma_), and neurite volume fraction (f_neurite_) from SANDI. The targets were the corresponding LSM-derived NeuN density and GFAP SAF maps. 80% of the brain voxels were randomly selected for model training, while the remaining 20% were held out for testing. Model performance was evaluated on a voxel-wise basis within each strain by calculating the correlation between predictions for the held-out voxels and the corresponding LSM-derived measurements. Random-forest regression models (200 trees) were implemented using scikit-learn (Pedregosa et al., 2018), and the relative importance of each dMRI feature was estimated using the mean decrease in impurity (scikit-learn’s default feature_importances_).

We explored how well macroscopic MRI-based models captured broad, coarse-grained variation in microscopic cellular features across the eight animals, by comparing mean MRI-based predictions of neuronal cell density with corresponding mean LSM-derived measurements for individual brains. This was performed in two ways: i) The RF models with SANDI and DKI metrics trained independently for each animal were used to obtain the mean MRI-based predictions across GM on held-out voxels. ii) Using leave-one-animal-out validation. For each fold, voxel-wise dMRI features and corresponding LSM-derived neuronal-density measurements from seven animals were pooled to train a random-forest regression model. The trained model was then applied to dMRI features from the eighth animal, which was entirely excluded from training. This was repeated eight times, such that each animal served once as the held-out test case. For each held-out animal, predicted neuronal density was averaged across grey matter and compared with the corresponding mean LSM-derived neuronal density. To ensure balance in training, the same number of voxels from the seven training animals was used for model fitting.

## Ethics Declaration

All procedures were conducted under license number PP4211803, granted by the UK Home Office under the Animals (Scientific Procedures) Act 1986, and approved by the University of Oxford’s ACER (Animal Care and Ethical Review) committee (22 October 2024).

## Data and code availability

Code for processing dMRI and LSM data, as well as for cross-modal registrations is available on: https://github.com/SPMIC-UoN/LSM-MRI-bioimaging.

Data, including multimodal MRI and whole-brain light-sheet microscopy in the same mice, will be made available to the community upon publication.

## Acknowledgements

This work was supported by Wellcome Trust’s Bioimaging Technology Development Awards (313367/Z/24/Z, 340940/Z/25/Z) and the European Research Council (ERC Consolidator 101000969). We also acknowledge support by the Medical Research Council (MRC IMPACT DTP MR/N013913/1) and the Biotechnology and Biological Sciences Research Council (BBSRC ALERT BB/Z515760/1). The authors would like to thank Dr Jana Koth from the MRC WIMM Wolfson Imaging Centre in Oxford for facilitating imaging experiments with the local Ultramicroscope BLAZE; as well as Prof Davide Calebiro and Dr Joao Correia from the COMPARE Advanced Imaging Facility at the University of Birmingham for assistance with acquisitions using the Ultramicroscope II system.

## Author Contributions

JMH: Conceptualization, Data curation, Formal analysis, Investigation, Visualization, Methodology, Writing – original draft, Writing – review and editing

FDA, SA, JPMP (equal contribution): Software, Data curation, Formal analysis, Methodology, Writing – original draft, Writing – review and editing

SCW, MT: Resources, Writing – review and editing

QS, MC: Software, Methodology, Writing – review and editing

CL, GJH, ER, KIC: Resources, Writing – review and editing

TDF: Investigation, Methodology, Resources, Writing – review and editing

AJ, AHS, ZM: Resources, Writing – review and editing

CL, CT, SJ, AFDH, KLM: Resources, Investigation, Methodology, Writing – review and editing

EG, JPL, SNS (equal contribution): Conceptualization, Resources, Investigation, Methodology, Supervision, Funding acquisition, Writing – original draft, Writing – review and editing.

