## Supplementary Material for "High-throughput light-sheet 3D microscopy of whole rodent brain microstructure for integration with MRI"

### Supplementary Materials


Jenna M. Hanmer[1], Francesco D'Antonio[1*], Stephania Assimopoulos[1*], Jose-Pedro Manzano-Patron[1*], Sydney C. Wood[2], Mohamed Tachrount[2], Qijia Shen[2], Martin Craig[1], Cyril Lai[3], Graham J. Hickman[4], Edalat Radfar[5], Kim I. Chisholm[6], Tracy D. Farr[7], Alan Jasanoff[8], Anna Hoerder-Suabedissen[9], Zoltan Molnar[9], Clemence Ligneul[2,10], Cristiana Tisca[2], Saad Jbabdi[2], Amy F.D. Howard[2,11], Karla L. Miller[2], Elizabeth de Guzman[2,*], Jason P. Lerch[2,12,13,*], Stamatios N. Sotiropoulos[1,*#]

[1]Sir Peter Mansfield Imaging Centre, School of Medicine, University of Nottingham, UK
[2]Oxford Centre for Integrative Neuroimaging, University of Oxford, UK
[3]MRC Weatherall Institute of Molecular Medicine, University of Oxford, UK
[4]School of Science & Technology, Nottingham Trent University, UK
[5]COMPARE Advanced Imaging Facility, College of Medicine and Health, University of Birmingham, UK
[6]School of Life Sciences, University of Nottingham, UK
[7]Institute for Neuroscience and Cardiovascular Research, University of Edinburgh, UK
[8]Department of Biological Engineering, Massachusetts Institute of Technology, Cambridge, MA, USA
[9]Department of Physiology, Anatomy and Genetics, University of Oxford, UK
[10]Molecular Imaging Research Center, Université Paris Saclay, CEA, Fontenay-aux-Roses, France
[11]Faculty of Engineering, Imperial College London, UK
[12]Department of Medical Biophysics, University of Toronto, Toronto, Canada
[13]Mouse Imaging Centre, The Hospital for Sick Children, Toronto, Canada
[*]These authors contributed equally - [#]Corresponding author

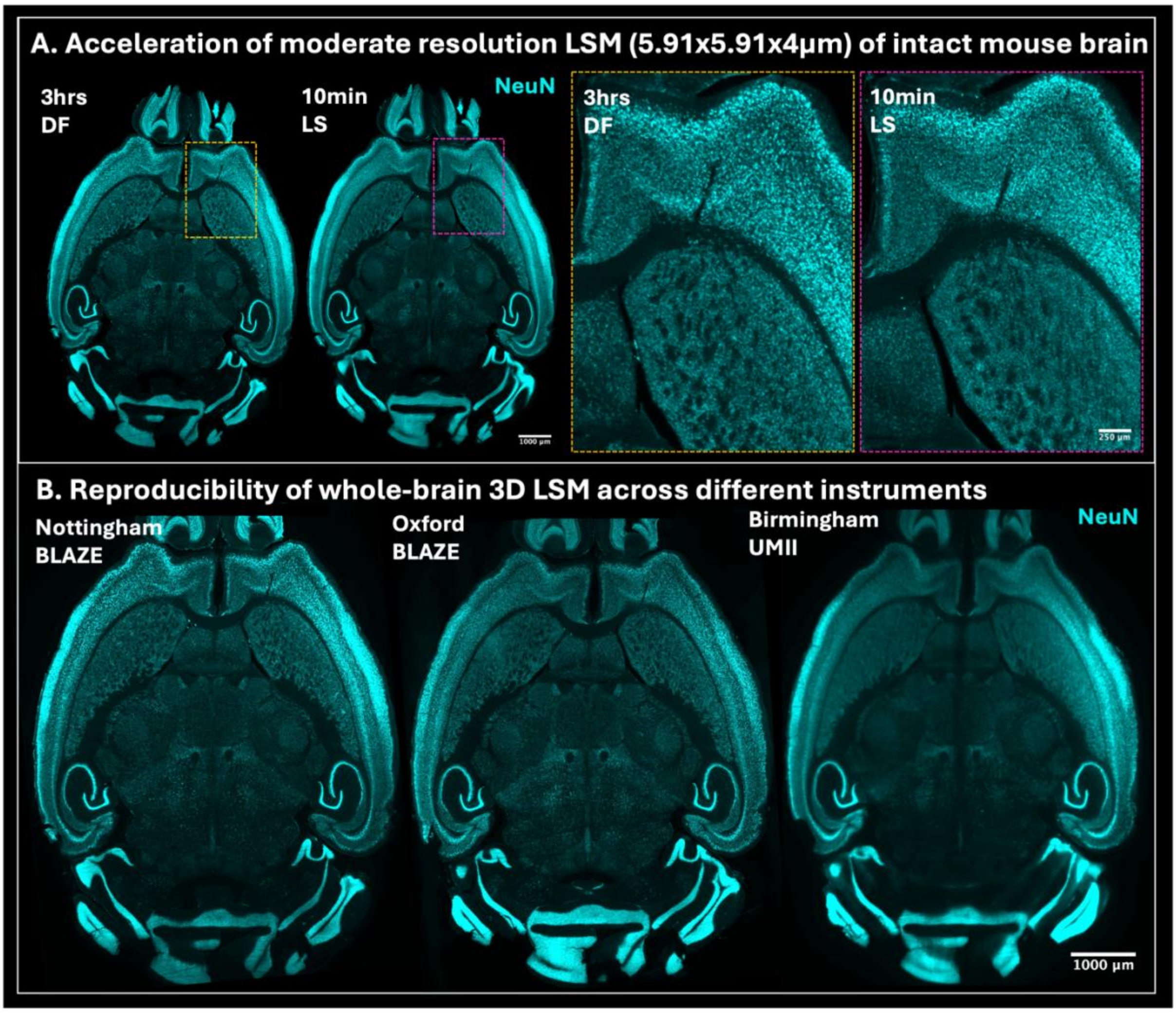


**SUPPLEMENTARY FIGURE 1. (A)** Comparison of conventional dynamic horizontal focus (DF; 13 steps) and light speed (LS) imaging using comparable acquisition parameters on the same instrument. Enlarged regions show that LS mode substantially accelerates acquisition without compromising image quality. **(B)** Reproducibility of DF imaging across three light-sheet microscopy systems. The same sample was imaged on an Ultramicroscope BLAZE in Nottingham and Oxford and an Ultramicroscope II in Birmingham, with all acquisitions completed within 3hrs. Comparable imaging performance across instruments and facilities demonstrates the generalisability of the methodology across systems with differences in hardware (e.g., cameras and optics) and software (e.g., file formats).

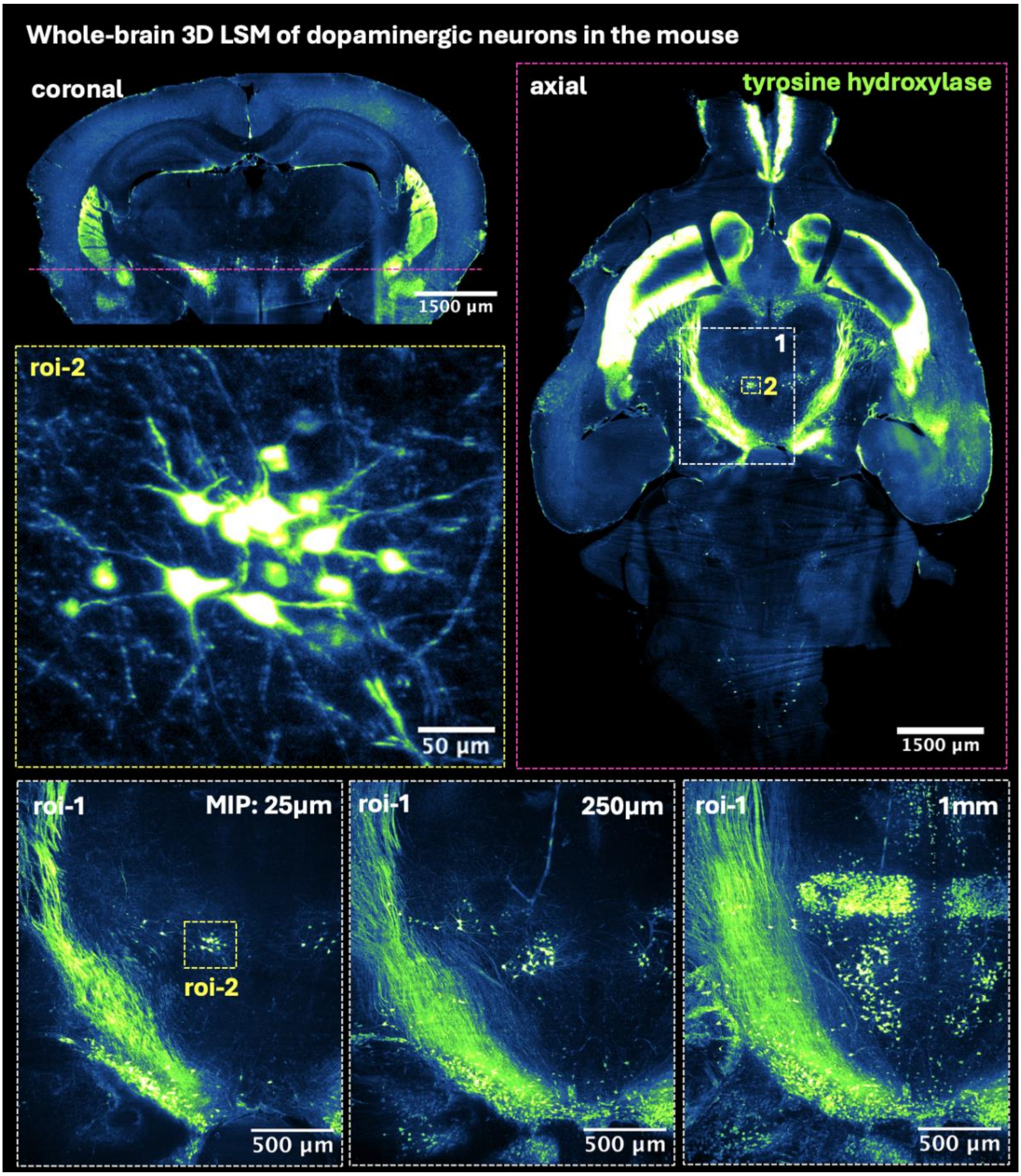


**SUPPLEMENTARY FIGURE 2.** Mouse brain labelled for tyrosine hydroxylase to visualise dopaminergic neurons and projections. **Top left:** Coronal view of the whole brain, with the purple line indicating the approximate position of axial slice shown on right. **Top right:** Axial view of the whole brain showing the approximate locations of regions 1 and 2. **Middle left:** Enlarged view of region 2, showing a cluster of dopaminergic neurons in the midbrain. **Bottom row:** Maximum intensity projection (MIP) of region 1, generated from image stacks of 25µm, 250 µm and 1mm thickness.

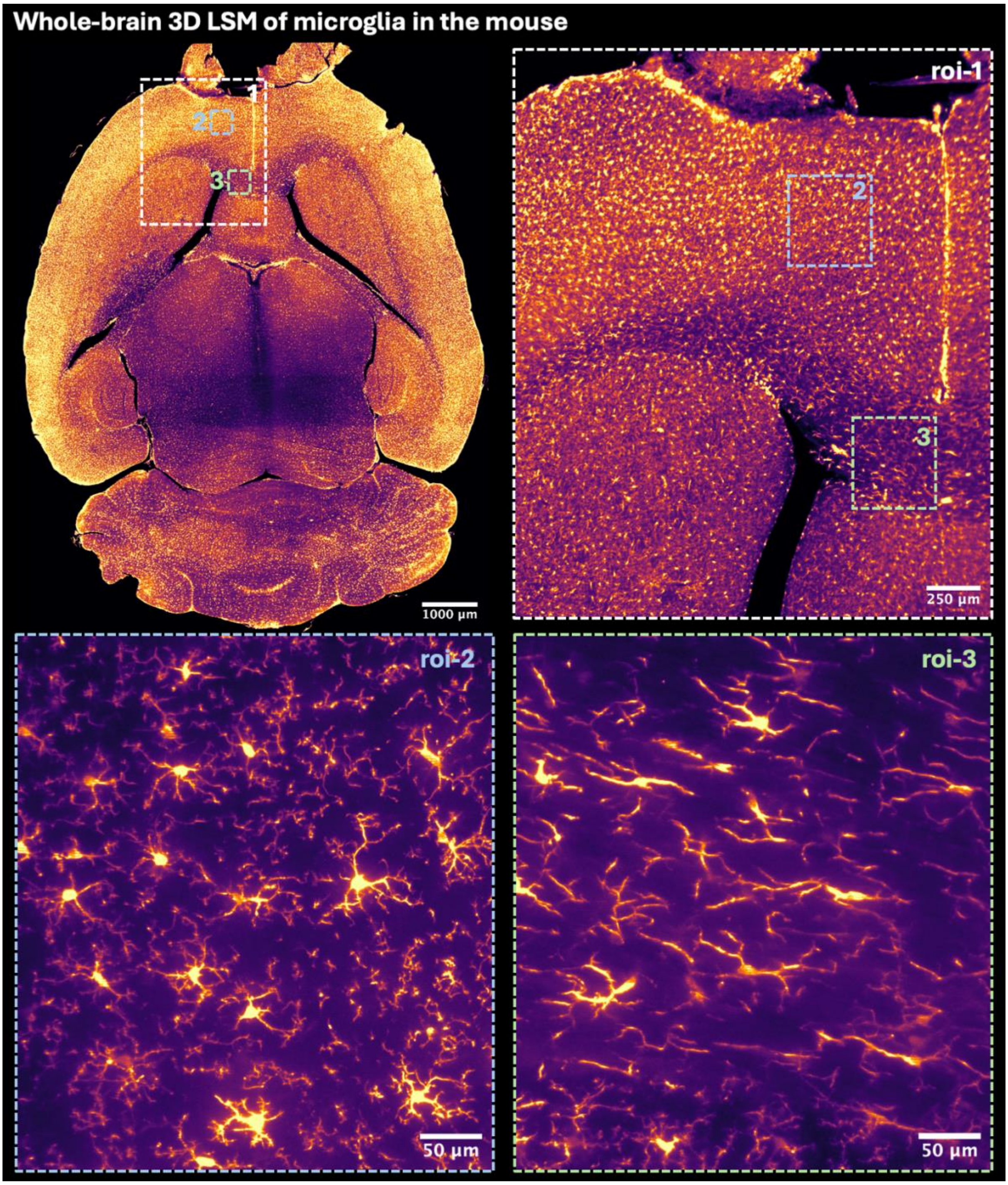


**SUPPLEMENTARY FIGURE 3.** Mouse brain labelled with IBA-1 (ionized calcium-binding adaptor molecule 1) to visualise microglia. **Top left:** axial slice of whole brain with approximate locations of regions 1, 2 and 3. **Top right:** Enlarged view of region 1. **Bottom row:** Enlarged views of regions 2 and 3, illustrating the morphological diversity of IBA-1-positive microglia.

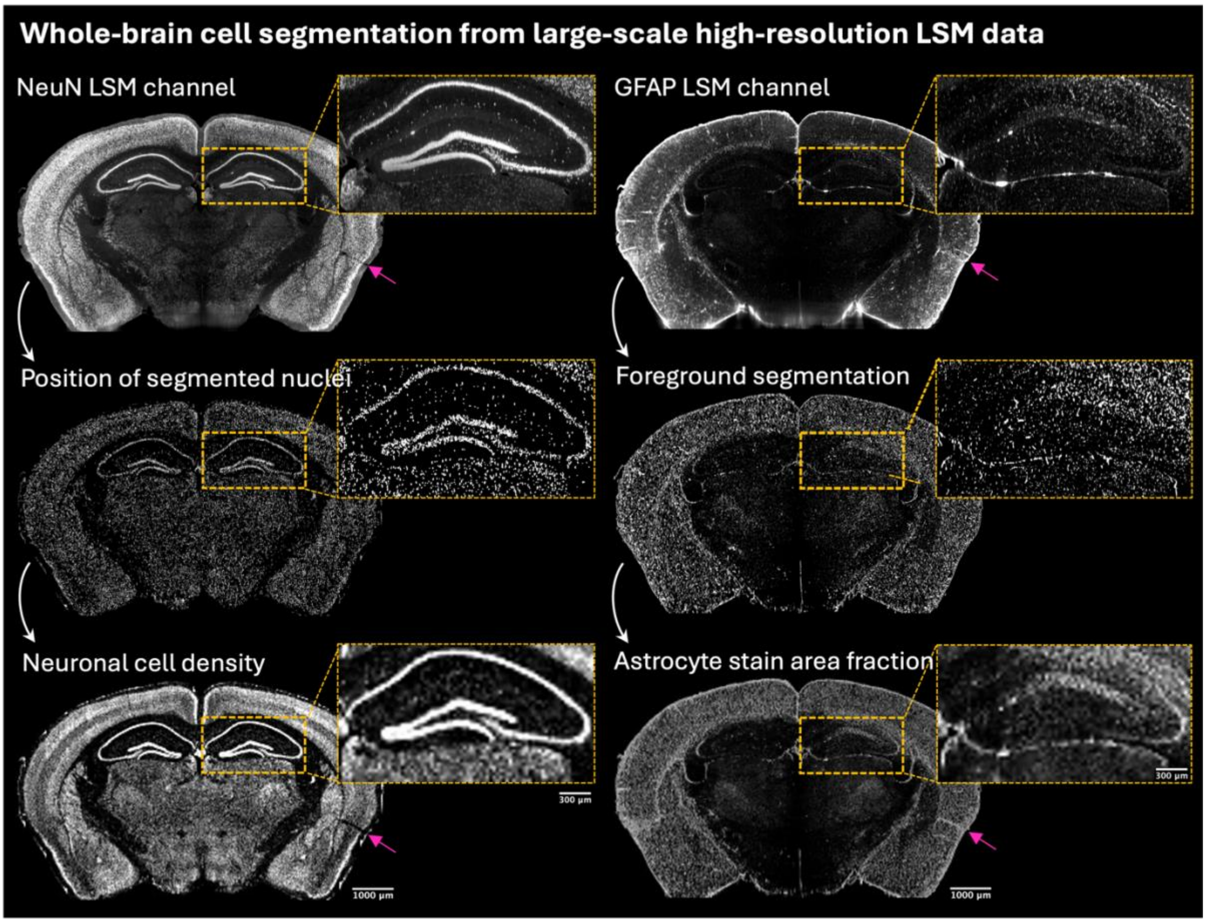


**SUPPLEMENTARY FIGURE 4.** Segmentation of neuronal nuclei and astrocyte signal from dual labelled whole mouse brain LSM. Left: NeuN-based neuronal cell counting, Right: GFAP stain area fraction (SAF). **Top row:** LSM fluorescence channels. **Middle row:** Segmentation outputs. **Bottom row:** resulting NeuN cell density and GFAP SAF maps. Pink arrows indicate preservation of left–right anatomical orientation throughout the processing pipeline.

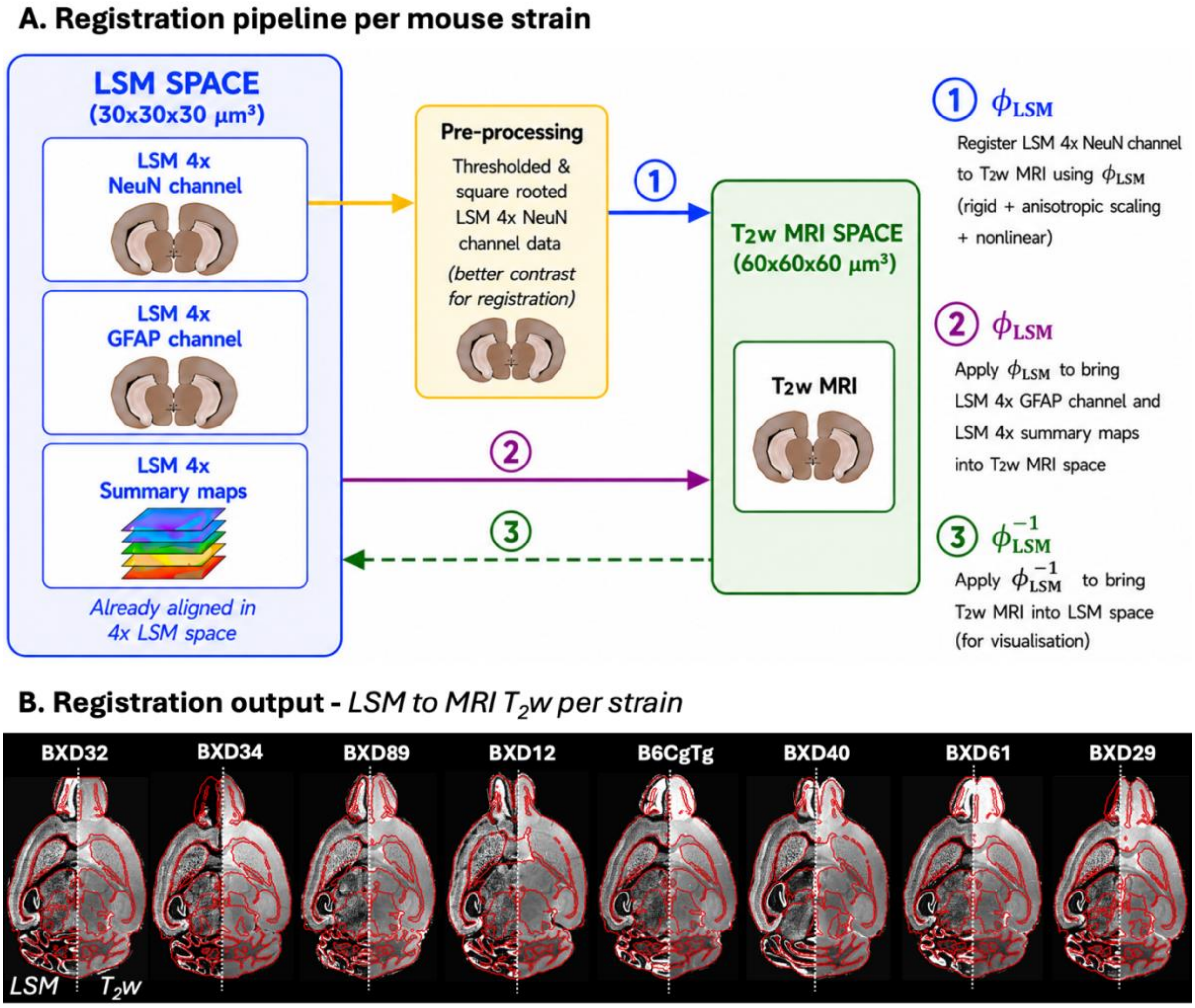


**SUPPLEMENTARY FIGURE 5.** Registration pipeline for aligning LSM and MRI data of the same mouse brain. **(A)** NeuN-labelled LSM images were thresholded to remove background and square-root transformed before non-linear registration to the corresponding $T_2$w MRI using TIRL (Huszar et al., 2023). The calculated forward transformation ($\phi_{LSM}$) was applied to NeuN- and GFAP-labelled LSM data, as well as the neuron density and astrocyte stain area fraction maps, to align these contrasts with the corresponding $T_2$w MRI. **(B)** Representative output of registration pipeline in LSM space, showing alignment between neuron density maps (left) and $T_2$w MRIs (right) across the mouse strains. A $T_2$w-derived grey matter mask is shown as a red overlay.

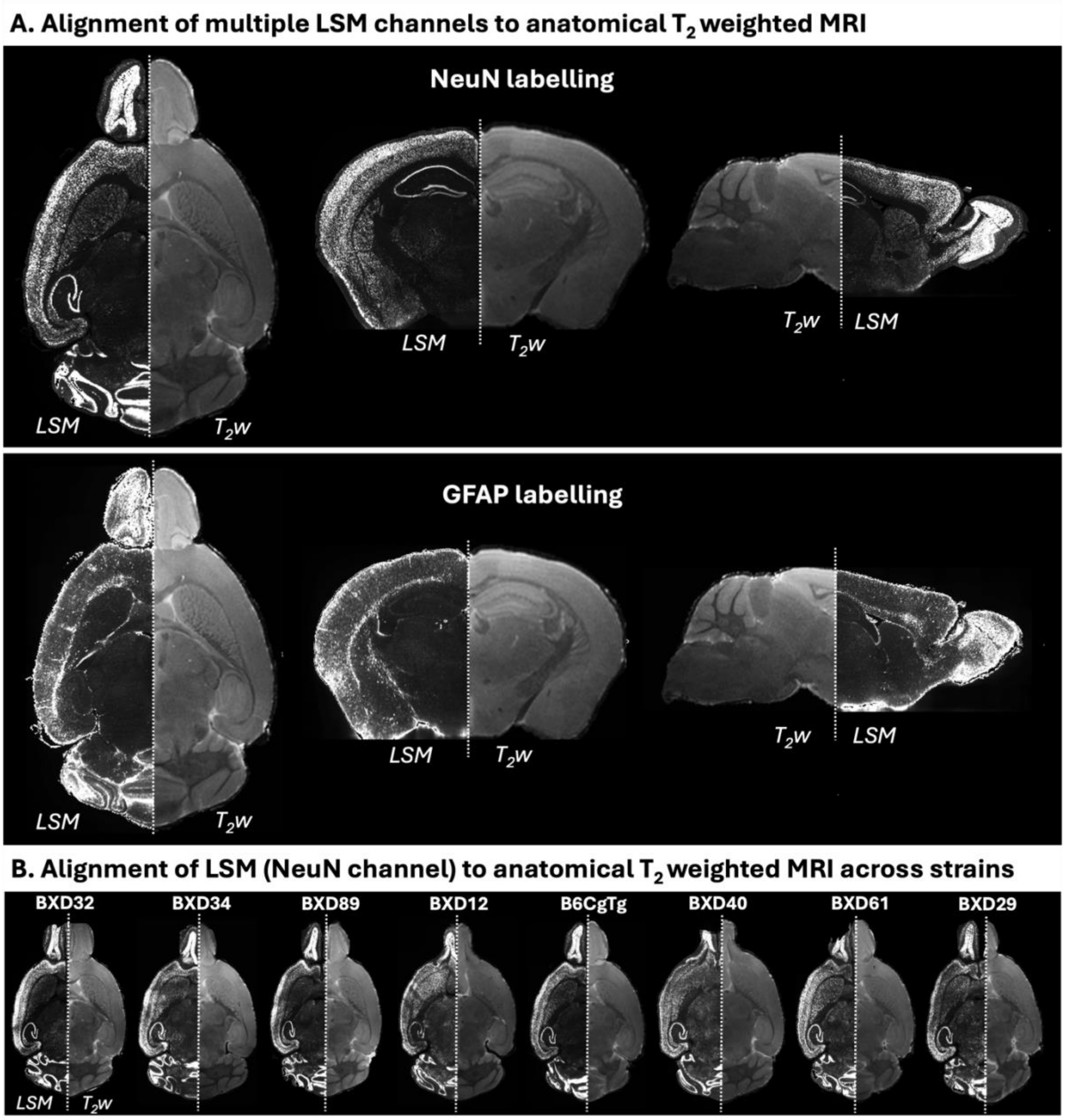


**SUPPLEMENTARY FIGURE 6.** Registration results showing alignment of the different LSM data types with anatomical $T_2$ weighted ($T_2$w) MRI. **(A)** Alignment of NeuN and GFAP labelling with $T_2$w MRI for one animal shown in the axial (LSM-left, $T_2$w-right), coronal (LSM-left, $T_2$w-right), and sagittal (LSM-right, $T_2$w-left) views. **(B)** Alignment of NeuN-labelled LSM data with $T_2$w MRI data across all mouse strains shown in the axial view (LSM-left, $T_2$w-right).

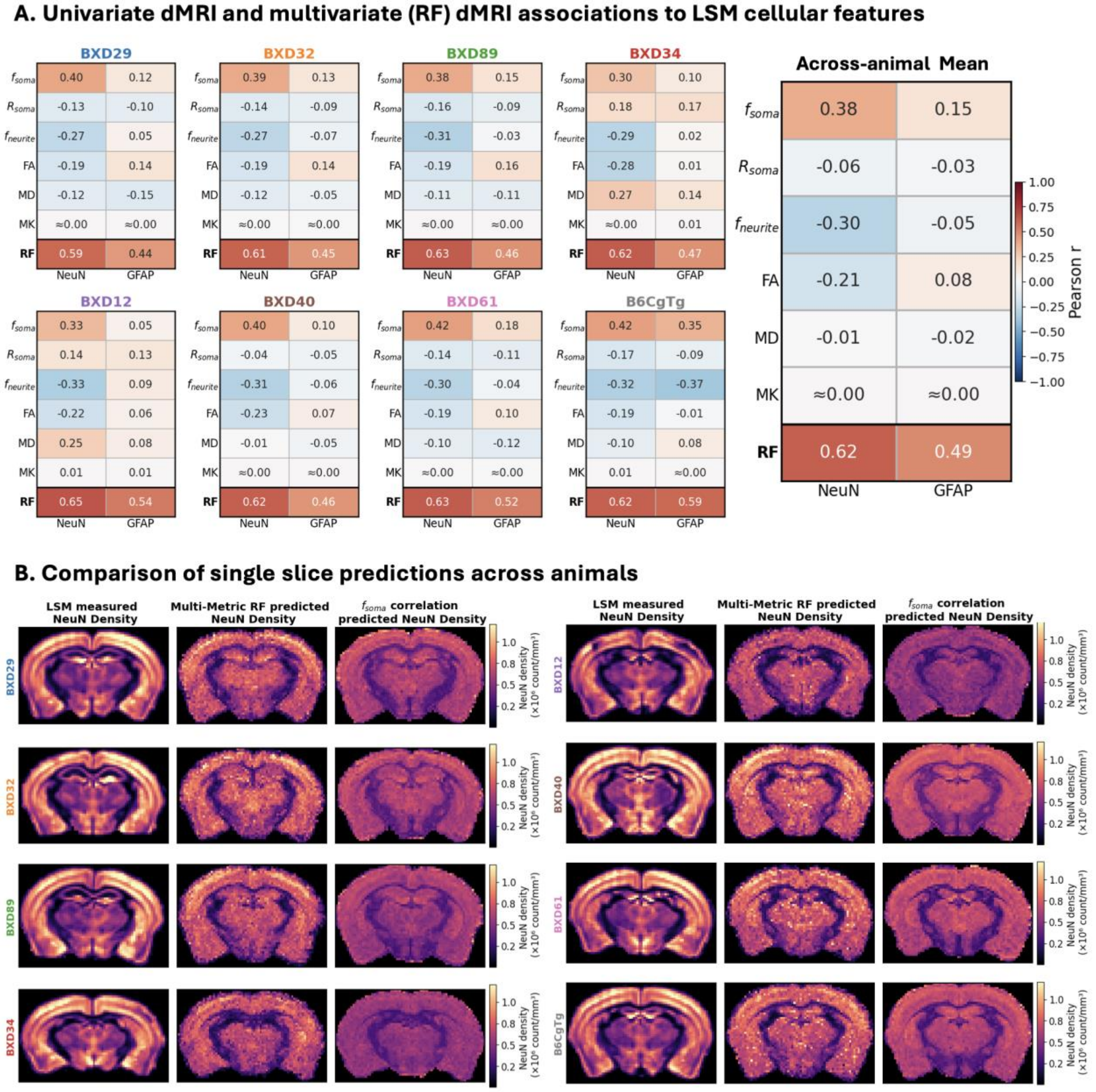


**SUPPLEMENTARY FIGURE 7. (A)** Individual SANDI and DKI metrics showed weaker associations with neuron cell density and GFAP stain area fraction than the predictive performance achieved by the multivariate random forest (RF) models combining all SANDI and DKI metrics. **(B)** LSM-derived neuronal cell density maps for all eight animals, alongside corresponding maps predicted by an RF model (held-out slice) incorporating all SANDI and DKI metrics and by a model using only the SANDI soma signal fraction ($f_{soma}$) as input.

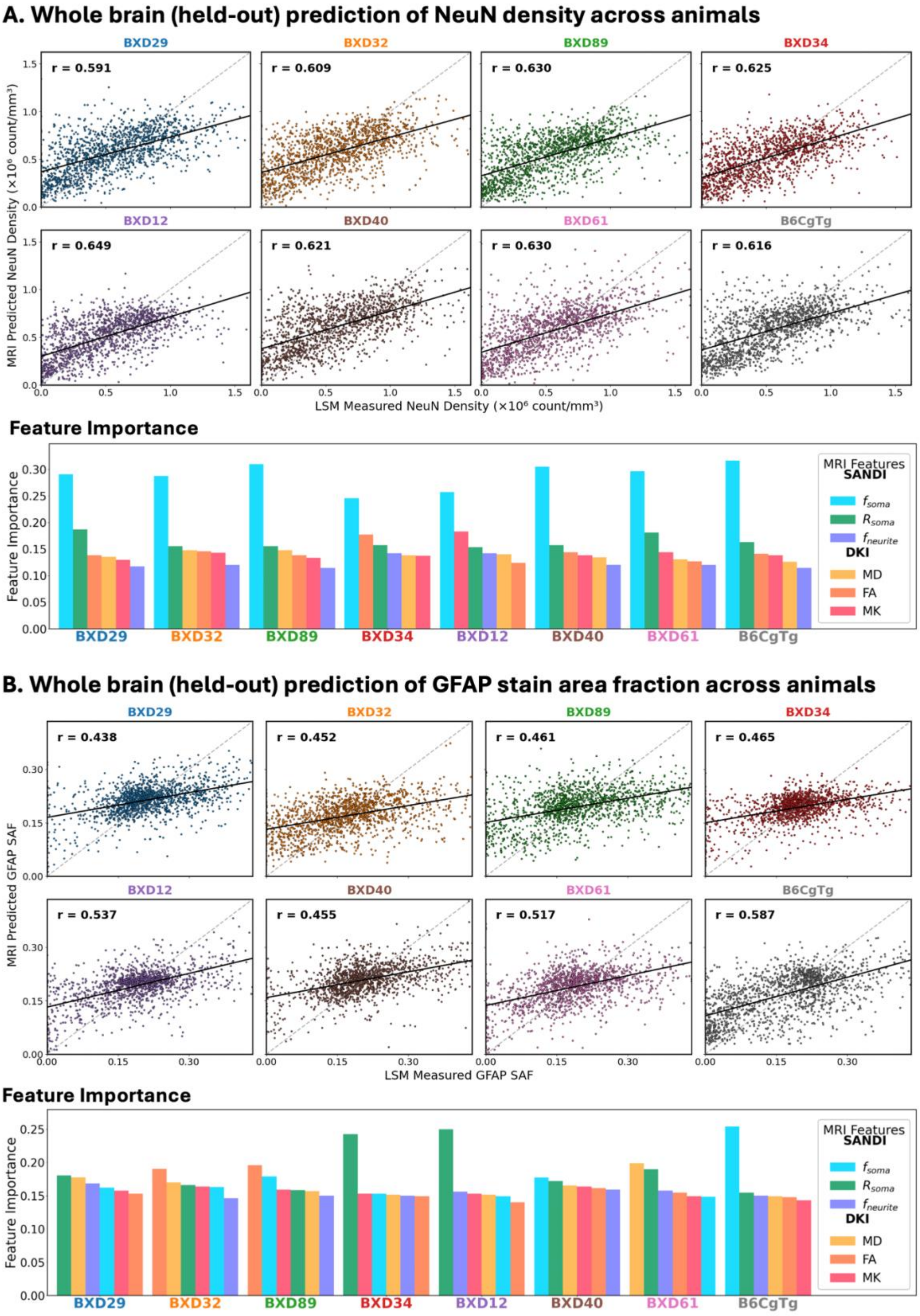


**SUPPLEMENTARY FIGURE 8. (A)** Neuron cell density predictions from MRI contrasts, assessed using held- out data, and relative importance of features (SANDI and DKI) across all eight animals. **(B)** Astrocyte (GFAP) stain area fraction (SAF) predictions from MRI contrasts, assessed using held out data, and relative importance of features (SANDI and DKI) across all eight animals.

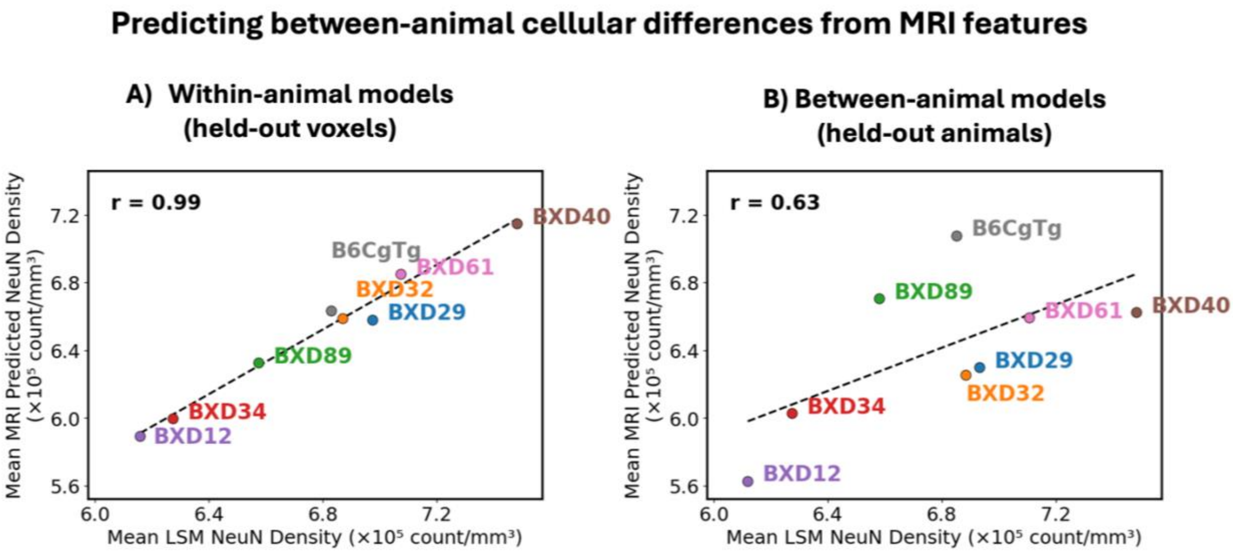


**SUPPLEMENTARY FIGURE 9.** Using dMRI features to predict between-animal cellular differences, as measured by LSM. MRI predictions averaged across grey matter are plotted against grey matter average LSM-derived neuronal density. Two types of models have been used: **(A)** Within-animal models. For each animal, an individual RF model was trained using 80% of voxels and predictions were obtained on 20% of held out voxels. Averages were obtained in held-out voxels. **(B)** Between-animal models. Each animal was left out of training and was predicted using an RF model trained on pooled dMRI-LSM data from the other seven animals.
